# Intestinal-derived FGF15 preserves muscle and bone mass following sleeve gastrectomy

**DOI:** 10.1101/2020.06.02.130278

**Authors:** Nadejda Bozadjieva Kramer, Jae Hoon Shin, Yikai Shao, Ruth Gutierrez-Aguilar, Ziru Li, Kristy M. Heppner, Samuel Chiang, Sara G. Vargo, Katrina Granger, Darleen A. Sandoval, Ormond A MacDougald, Randy J. Seeley

## Abstract

Bariatric surgeries such as the Vertical Sleeve Gastrectomy (VSG) are invasive, but provide the most effective long-term metabolic improvements in obese and Type 2 diabetic patients. These powerful effects of manipulating the gastrointestinal tract point to an important role of gastrointestinal signals in regulating both energy balance and metabolism. To that end, we have used mouse models of VSG to identify key gut signals that mediate these beneficial effects. Preliminary data from our rodent model of VSG led us to hypothesize a potential role for the hormone Fibroblast-Growth Factor15/19 (mouse/human ortholog) which pharmacologically can regulate many aspects of energy homeostasis and glucose handling. FGF15 is expressed in ileal enterocytes of the small intestine and is released postprandially. Like many other gut hormones, postprandial plasma levels in humans and ileal FGF15 expression in mice increase after VSG. We generated intestinal-specific FGF15 knock out (VilCreERT2; Fgf15^f/f^) mice and controls, which were maintained on 60% high-fat diet. Interestingly, ablation of intestinal FGF15 in adult mice results in little change to body weight or glucose regulation when challenged with a high-fat diet. Unlike what we had predicted, intestinal-specific FGF15 knock out mice lost more weight after VSG and this was a result of increased lean tissue loss compared to control mice. Further, the loss of bone mineral density observed after VSG in control mice was increased in intestinal-specific FGF15 knock out mice. Finally the effect of VSG to reduce hepatic cholesterol was also absent in intestinal-specific FGF15 knock out mice. These data point to an important role for intestinal FGF15 to protect the organism from deleterious effects of rapid weight loss that occurs after VSG.

## Introduction

Obesity has become a growing epidemic, where associated complications such as cardiovascular morbidity, type 2 diabetes and insulin resistance pose major health care challenges worldwide (Afshin et al., 2017). Current pharmacological treatments for obesity include less than ten FDA approved drugs, all of which demonstrate modest effect sizes and substantial liabilities for patients including cardiac and gastrointestinal distress (Saltiel, 2016). Although invasive, bariatric surgery is the most effective treatment for sustained weight loss, and also improves glycemic control and other comorbidities in patients better than conventional weight-loss therapies (Adams et al., 2017; Schauer et al., 2017).

The effectiveness of bariatric surgery to reduce body weight and improve glucose metabolism has highlighted the important role that the gastrointestinal tract has in regulating a wide range of metabolic processes. One weight-independent effect of bariatric surgery is the alteration of enterohepatic bile acid circulation resulting in increased plasma bile levels as well as altered bile acid composition in rodents (Kohli et al., 2010; Myronovych et al., 2014) and humans (Patti et al., 2009; Pournaras et al., 2012). While it remains unclear why both VSG and RYGB can alter bile acids, it is possible that these changes are important mediators of the effects of surgery. We have previously identified bile acid signaling through the nuclear ligand-activated farnesoid X receptor (FXR) as a potential link for mediating the beneficial effects of elevated bile acids following bariatric surgery. We have shown that bile acids are increased after VSG and that FXR is essential for the positive effects of bariatric surgery on weight loss and glycemic control (Myronovych et al., 2014; Ryan et al., 2014). Unlike wild-type mice, *FXR−/−* mice do not maintain body weight loss and do not have improved glucose tolerance after VSG or after bile diversion to the ileum (Albaugh et al., 2019; Ryan et al., 2014). These studies have highlighted the importance of the enterohepatic circulation in the metabolic effects following bariatric surgery.

Downstream of FXR is the gut-derived hormone Fibroblast Growth Factor 15 (the human ortholog is termed FGF19). Pharmacological administration of FGF15/19 has potent effects to reduce bile acid secretion at the level of both the liver and the gallbladder (Kir et al., 2011; Potthoff et al., 2011) and has potent effects on body weight and glucose maintenance (Lan et al., 2017). In the ileum, bile acids activate intestinal FXR and its downstream target FGF15/19. FGF15/19 is expressed in ileal enterocytes of the small intestine and is released postprandially in response to bile acid absorption (Inagaki et al., 2005). Once released from the ileum, FGF15/19 enters the portal venous circulation and travels to the liver where FGF15/19 binds to its receptor FGFR4 and represses de novo bile acid synthesis and gallbladder filling (Inagaki et al., 2005). Therefore, bile acids and FGF15/19 both act as ligands to regulate bile acid synthesis and facilitate communication between the liver and small intestine. The actions of FGF15/19 resemble that of insulin in stimulating protein and glycogen synthesis and reducing gluconeogenesis. However, unlike insulin, FGF15/19 decreases hepatic triglycerides and reduces cholesterol. This notable difference has made FGF19 a potential therapeutic target to aid insulin’s actions while avoiding some pitfalls of insulin therapy (DePaoli et al., 2019; Harrison et al., 2018).

Consistent reports demonstrate that pharmacologically elevating Fibroblast Growth Factor FGF15/19 levels in preclinical models of metabolic disease results in multiple metabolic benefits including increased energy expenditure, reduced adiposity, and improved lipid and glucose homeostasis (Fu et al., 2004; Miyata et al., 2011; Morton et al., 2013; Ryan et al., 2013; Tomlinson et al., 2002). Circulating FGF19 levels are reduced in individuals with metabolic disorders and nonalcoholic fatty liver disease (NAFLD). FGF19 levels are lower in obese patients, without strong association to glucose metabolism or insulin sensitivity (Gallego-Escuredo et al., 2015; Gomez-Ambrosi et al., 2017; Haluzikova et al., 2013; Mraz et al., 2011; Renner et al., 2014). Other studies report that basal FGF19 levels are inversely correlated to glucose metabolism or insulin sensitivity (Barutcuoglu et al., 2011; Sonne et al., 2016) and nonalcoholic fatty liver disease (NAFLD) (Eren et al., 2012; Jiao et al., 2018). Most importantly, fasting and postprandial plasma FGF19 levels in humans (Belgaumkar et al., 2016; DePaoli et al., 2019; Gomez-Ambrosi et al., 2017; Haluzikova et al., 2013; Mulla et al., 2019; Sachdev et al., 2016) and ileal FGF15 expression in mice (as shown in these studies) increase after VSG. These data point to FGF15/19 as a potential target to mediate the effects of weight loss and improved glucose tolerance following VSG. To test the hypothesis of whether FGF15 plays a role in the metabolic improvements after bariatric surgery, we used a mouse model of VSG, which has increased ileal expression of FGF15 in VSG compared to Sham mice. We generated a novel mouse model of intestinal-specific FGF15 knock out (VilCreERT2; Fgf15^f/f^) and controls, which were maintained on 60% high-fat diet before and after undergoing Sham or VSG surgery. Intestinal-derived FGF15 is necessary for the improvement in peripheral blood glucose regulation, to preserve muscle mass and bone mass, and for the decrease in hepatic cholesterol after VSG-induced weight loss. These finding point to an important role for FGF15 in the regulation of multiple metabolic parameters following VSG.

## Results

### Intestinal FGF15 expression increases and prevents muscle mass loss after VSG in mice

Circulating FGF19 levels are lower in obese patients (Gallego-Escuredo et al., 2015; Gomez-Ambrosi et al., 2017; Haluzikova et al., 2013; Mraz et al., 2011; Renner et al., 2014) and patients with impaired glucose metabolism or insulin sensitivity (Barutcuoglu et al., 2011; Sonne et al., 2016) and nonalcoholic fatty liver disease (NAFLD) (Eren et al., 2012; Jiao et al., 2018). This highlights the relationship of FGF19 with body weight and metabolism. Numerous reports have also shown that plasma FGF19 levels increase after weight-loss surgeries (Belgaumkar et al., 2016; DePaoli et al., 2019; Gomez-Ambrosi et al., 2017; Haluzikova et al., 2013; Mulla et al., 2019; Sachdev et al., 2016). Due to the lack of commercially available and validated FGF15 assays, we were unable to measure circulating FGF15 levels after VSG in mice (Angelin et al., 2012; Montagnani et al., 2011). However, our data show that ileal FGF15 expression increases after VSG in mice (Figure 1A). These data point to FGF15/19 as a potential target to mediate the effects of weight-loss surgery.

**Figure 1.**
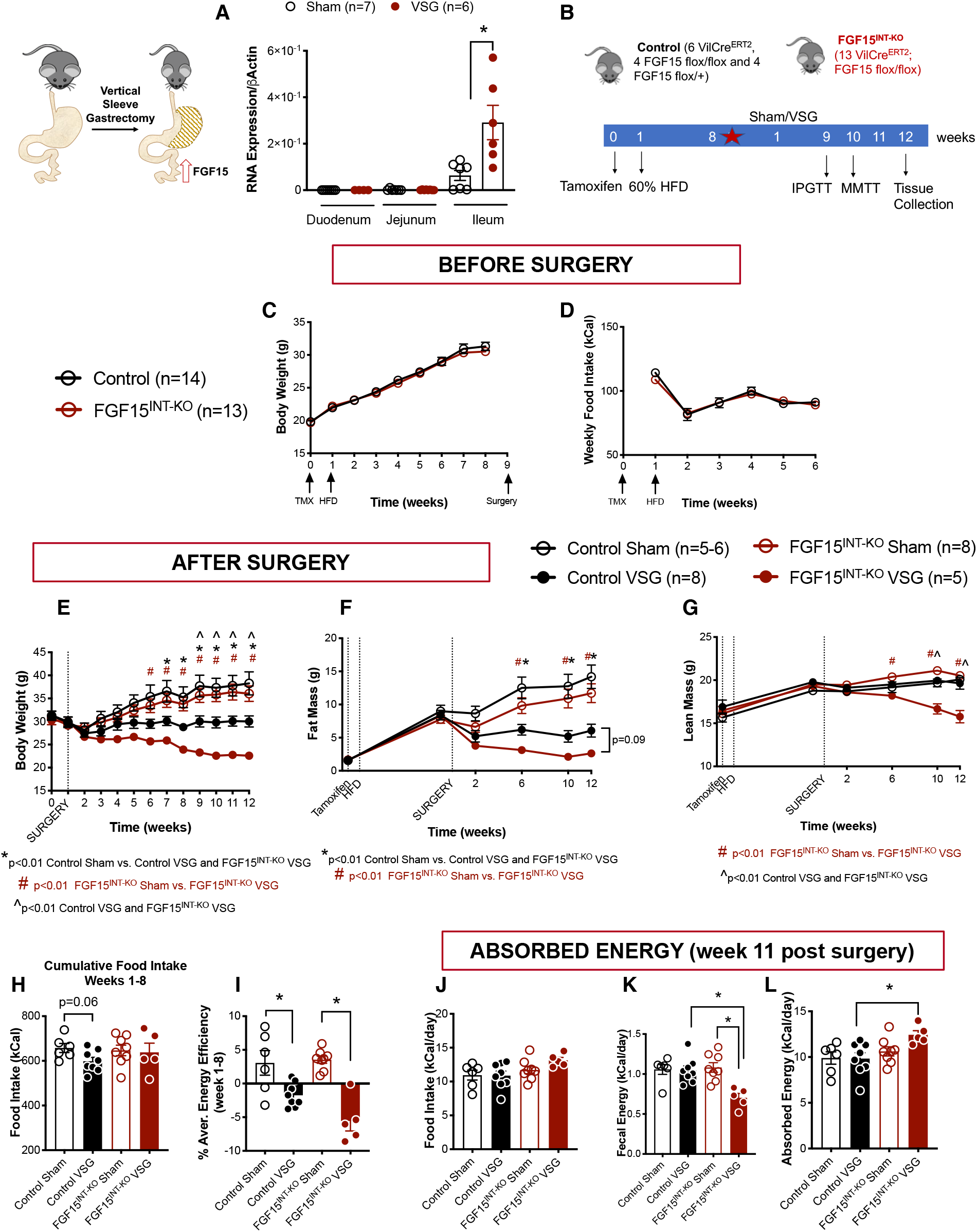
Intestinal FGF15 expression increases and prevents muscle mass loss after VSG in mice. **A.** FGF15 RNA expression in duodenum, jejunum and ileum in Sham and VSG mice fed 60% HFD. Intestinal mucosa was collected 15 minutes post mixed meal gavage (n=6-7). **B.** Experimental timeline. **C.** Body weight and **D.** Food intake before surgery. **E.** Body weight after surgery. **F.** Fat body mass and **G.** Lean body mass before and after surgery. **H.** Cumulative food intake and **I.** Energy Efficiency for weeks 1-8 after surgery. **J.** Food intake, **K.** Fecal energy and **L.** Absorbed energy during week 11 after surgery. Animal number WT Sham (n=5-6), WT VSG (n=8), FGF15^INT-KO^ Sham (n=8), FGF15^INT-KO^ VSG (n=5). Data are shown as means ± S.E.M. *p<0.01 Control Sham vs. Control VSG; #p<0.01 FGF15^INT-KO^ VSG and FGF15^INT-KO^ Sham; ^p<0.01 Control VSG and FGF15^INT-KO^ VSG. Panels A, C and D were analyzed with Student’s 2-tailed *t* test. Panels E-L were analyzed with 2-Way ANOVA with Tukey’s post-test).

Global ablation of FGF15 in FGF15−/− mice resulted in impaired glucose tolerance (Kir et al., 2011; Potthoff et al., 2011), but in our hands these mice are surprisingly protected against diet-induced obesity (data not shown). FGF15 is highly expressed in the developing mouse brain (Gimeno et al., 2003; Gimeno et al., 2002; McWhirter et al., 1997). We speculate that the total body knockouts have impaired development of the CNS, which likely contributes to their reduced weight gain on a high-fat diet. Expression of FGF15 in the adult mouse becomes limited to the distal intestine and dorsal medial hypothalamus (Fon Tacer et al., 2010; Inagaki et al., 2005; Picard et al., 2016). These issues make it difficult to use the total body knockout to determine critical aspects of FGF15 function in the adult animals. Due to this weight difference on HFD and great difficulty breeding whole body knockouts, we generated a novel FGF15 flox/flox mouse to test specific hypotheses about the tissue-specific role of FGF15 in metabolism after VSG. The FGF15 flox/flox mice, were built using CRISPR-Cas9 with LoxP sites flanking exon 2 of the FGF15 gene. We bred these mice to VilCreERT2 mice and administered tamoxifen (intraperitoneal, 3 doses/150 mg/kg; Figure 1B) to FGF15^INT-KO^ (VilCreERT2; FGF15 flox/flox) and Controls (6 VilCreERT2, 4 FGF15 flox/flox and 4 FGF15 flox/+ were combined for comparisons).

After the studies were completed, we validated exon 2 excision within ileal mucosa (where FGF15 is most highly expressed) in all mice (Supplemental Figure 1). All mice received tamoxifen at the same time, and a week later were placed on 60% high fat diet (HFD; Figure 1B). Prior to surgery, Control and FGF15^INT-KO^ mice had similar body weight increase in response to HFD, without differences in food intake, fat and lean mass (Figure 1C, D, F, G).

After 8 weeks of being on HFD, each mouse underwent either a Sham or VSG procedure and were returned to HFD four days after surgery. As expected, Control mice receiving VSG lost a significant amount of body weight compared to Control Sham mice, and that weight loss came mostly from loss of fat mass (Figure 1E, F). Control VSG mice maintained their lean mass even after surgery, as we have shown before (Patel et al., 2018). FGF15^INT-KO^ VSG mice also lost a significant amount of weight, but 9 weeks post-surgery their body weight was significantly lower compared to Control VSG (Figure 1E). Although FGF15^INT-KO^ VSG mice lost fat mass, surprisingly they also lost a significant amount of lean mass and this loss was significantly greater than when compared to Control VSG and both Sham groups (Figure 1F, G).

Cumulative food intake and the average energy efficiency (change in body weight divided by food intake) for the 8 weeks after surgery showed that the body weight loss observed in FGF15^INT-KO^ VSG mice was not a result of decreased food intake (Figure 1H, I). In fact, although Control VSG mice had reduced cumulative food intake for the 8 weeks post-surgery (with the largest difference observed in the first few weeks after surgery), FGF15^INT-KO^ VSG mice had no significant reduction in food intake (Figure 1H). This introduced the possibility that the loss of intestinal FGF15 leads to increased malabsorption following VSG. We measured the absorbed energy by bomb calorimetry 11 weeks after surgery. The average daily food intake was not different between the groups (Figure 1J). However, the fecal energy output was reduced in FGF15^INT-KO^ VSG mice compared to Control VSG and FGF15^INT-KO^ Sham mice (Figure 1K). Given that all mice were on the same diet, FGF15^INT-KO^ VSG mice had increased their absorption of calories from ingested food (Figure 1L).

### Loss in muscle mass is accompanied by decreased strength and skeletal muscle fiber size in FGF15^INT-KO^ VSG mice

Previous data has shown that exogenous administration or genetic overexpression of FGF19 decreases body weight and adiposity (Fu et al., 2004; Lan et al., 2017; Miyata et al., 2011; Morton et al., 2013; Ryan et al., 2013; Tomlinson et al., 2002), but prevents muscle mass wasting by enlarging muscle fiber size and protecting muscle from atrophy (Benoit et al., 2017). In addition to significant loss of muscle mass in FGF15^INT-KO^ VSG mice (Figure 1G), we observed a trend of decreased grip strength (9 weeks post-surgery) and soleus muscle fiber size in these mice (Supplemental Figure 2A, B). Moreover, a distribution analysis of the fiber size showed that FGF15^INT-KO^ VSG mice have a rightward shift towards an increased number of smaller, and a lower number of larger soleus fibers (Supplemental Figure 2C).

To further examine the atrophy-related pathways in soleus muscle, we measured circulating Myostatin (GDF8) and Activin A levels and the soleus muscle expression of their downstream targets, muscle-specific E3 ubiquitin ligases Atrogin 1 and MuRF1. We did not observe any differences in the plasma Myostatin and Activin A levels, nor the RNA expression of Atrogin 1 and MuRF1 in soleus muscle (Supplemental Figure 2D-G). Next, we examined plasma levels of IGF1 and FGF21 as anti and pro-atrophy mediators in skeletal muscle. Twelve weeks after surgery postprandial plasma levels of IGF1 were decreased in FGF15^INT-KO^ VSG mice compared to Control VSG (Supplemental Figure 2H). Studies have indicated that FGF21 levels increase with fasting and FGF21 is necessary for the fasting-induced muscle mass and force loss (Oost et al., 2019). FGF15^INT-KO^ VSG mice had increased plasma levels of FGF21 compared to Control VSG and FGF15^INT-KO^ Sham mice (Supplemental Figure 2I). Additionally, there was a significant correlation between muscle mass and plasma FGF21 levels (Supplemental Figure 2J).

FGF15^INT-KO^ VSG mice did not show differences in small bowel biometry (Supplemental Figure 3A-C). The large bowel weight was significantly less in FGF15^INT-KO^ VSG compared to FGF15^INT-KO^ Sham mice (Supplemental Figure 3D). However, there was also a trend toward decreased large bowel weight in Control VSG compared to Control Sham mice, suggesting that decrease in large bowel weight could be related to decreased body weight of the VSG groups (Supplemental Figure 3D) (Mao et al., 2013). The ratio of large bowel weight/length was significantly reduced in both VSG groups compared to Control Sham, but was not significantly different compared to FGF15^INT-KO^ Sham mice (Supplemental Figure 3F). Further analysis of ileum cross-sections showed no differences in villi height, crypt depth or the ratio of villi height/crypt depth (Supplemental Figure 3G-I).

### Intestinal-derived FGF15 partially preserves bone and bone marrow adipose tissue (BMAT) loss following VSG

Previous studies in our lab have shown that loss of bone and BMAT (bone marrow adipose tissue) following VSG is independent of body weight and diet (Li et al., 2019). In the current study, VSG caused a precipitous loss of trabecular and cortical bone loss in FGF15^INT-KO^ mice. Specifically, FGF15^INT-KO^ VSG mice had decreased trabecular bone volume fraction (Tb. BV/TV), trabecular bone mineral density (Tb. BMD), and trabecular connective density (Conn. Dens) (Figure 2A-D). Although the thickness of the trabecular bone (Tb. Th) was not altered, the trabecular number (Tb. N) was decreased, while spacing between trabeculae was increased (Tb. Sp) in FGF15^INT-KO^ VSG mice (Figure 2 E-G). Cortical thickness (Ct. Th), bone area (Ct. BA/TA) and bone mineral density (Ct. BMD) were reduced by VSG and further decreased by lack of intestinal-derived FGF15 (Figure 2H-K).

**Figure 2.**
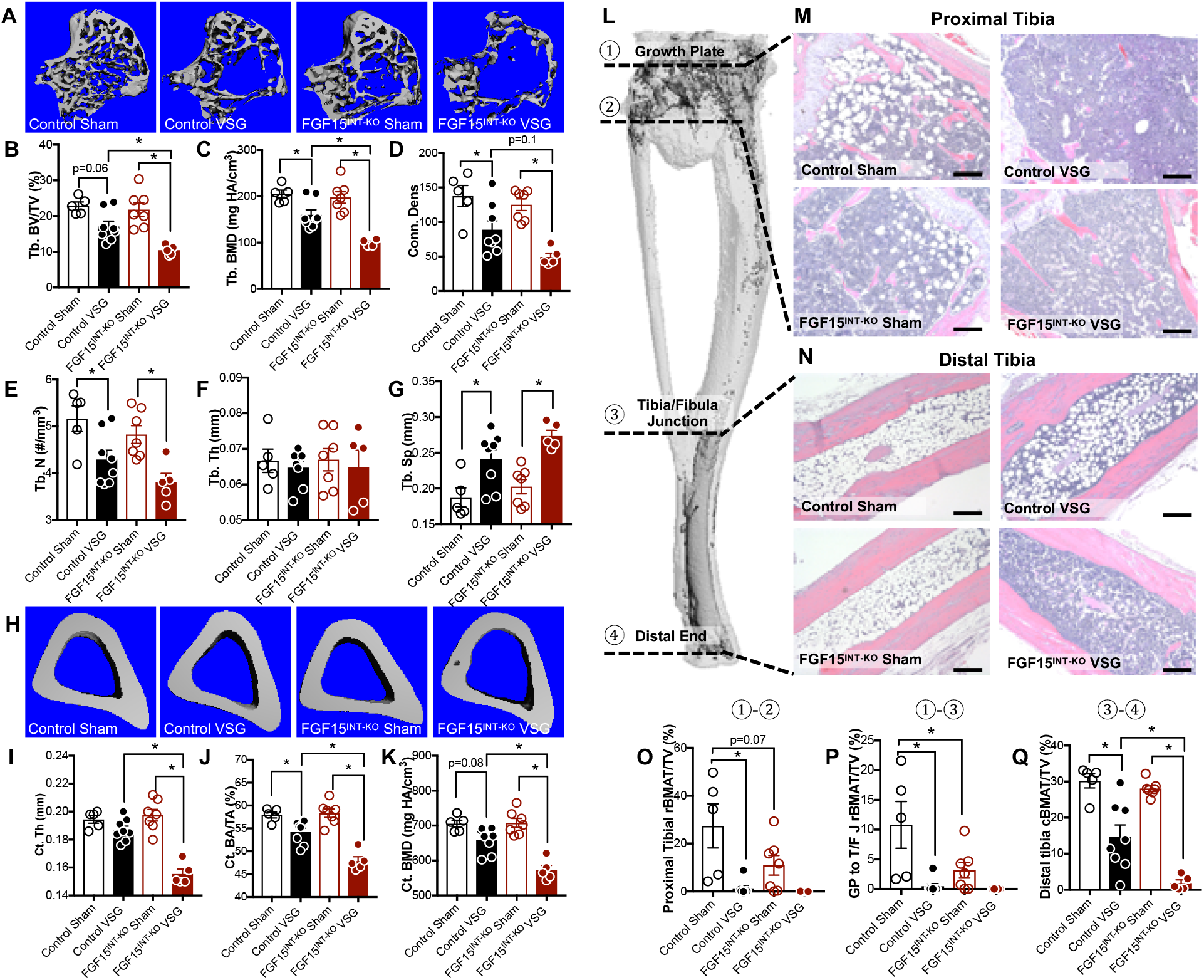
Intestinal-derived FGF15 partially preserves bone and BMAT loss following VSG. **A.** 3D images of trabecular bone. **B.** Trabecular bone volume fraction (Tb. BV/TV), **C.** Trabecular bone mineral density (Tb. BMD), **D.** Trabecular bone connective density (Conn. Des) and **E.** Trabecular number (Tb. N), **F.** Thickness of the trabecular bone (Tb. Th), **G.** Spacing between trabeculae (Tb. Sp). **H.**3D images of mid-cortical bone. **I.** Thickness of the cortical bone (Ct. Th). **J.** Cortical bone area (Ct. BA/TA) and **K.** Cortical bone mineral density (Ct. BMD). **L.** Tibial BMAT was visualized by osmium staining. **M-N.** Representative sections from proximal and distal tibiae were stained with H&E and are shown at ×100 magnification. Scale bars, 200 μm. **L.** Tibial BMAT was quantified relative to total bone volume after osmium staining within the indicated regions as shown in **O.** Proximal Tibia (①-②), **P.** Growth plate (G/P) to tibia/fibula junction (T/FJ) (①-③), **Q.** Distal tibia is T/F J to distal end (③-④). Animal number Control Sham (n=5), Control VSG (n=8), FGF15^INT-KO^ Sham (n=8), FGF15^INT-KO^ VSG (n=5). Data are shown as means ± S.E.M. * p<0.05 (2-Way ANOVA with Tukey’s post-test).

Consistent with our previous studies, both “regulated” bone marrow adipose tissue, BMAT (rBMAT) in proximal tibia and “constitutive” BMAT (cBMAT) in distal tibia were decreased by VSG (representative images Figure 2M, N) (Li et al., 2019; Scheller et al., 2015). Interestingly, loss of intestinal FGF15 caused a baseline reduction of rBMAT in proximal tibia ranging from growth plate (GP) to tibia/fibula junction (T/F J), with nearly complete depletion after VSG (Figure 2M, O, P). Although the distal tibial cBMAT of Sham mice was unaffected by loss of intestinal FGF15, this BMAT depot was more thoroughly depleted in FGF15^INT-KO^ VSG mice compared to Control VSG mice (Figure 2N, Q).

### Loss of intestinal FGF15 increases circulating total GLP-1 levels and increases gastric emptying after VSG

Four-hour fasted blood glucose concentrations were not different among any of the four groups, but insulin concentrations were lower in FGF15^INT-KO^ VSG compared to FGF15^INT-KO^ Sham (Figure 3A, B). Six and nine weeks after surgery, intraperitoneal glucose tolerance was improved in Control VSG mice as compared to their sham controls (Figure 3C, D). As expected, we saw a significant improvement in glucose tolerance in Control VSG compared to Control Sham mice six and nine weeks post-surgery (Figure 3C, D). To our surprise, there was no difference in glucose tolerance between FGF15^INT-KO^ VSG and FGF15^INT-KO^ Sham mice (Figure 3C, D).

**Figure 3.**
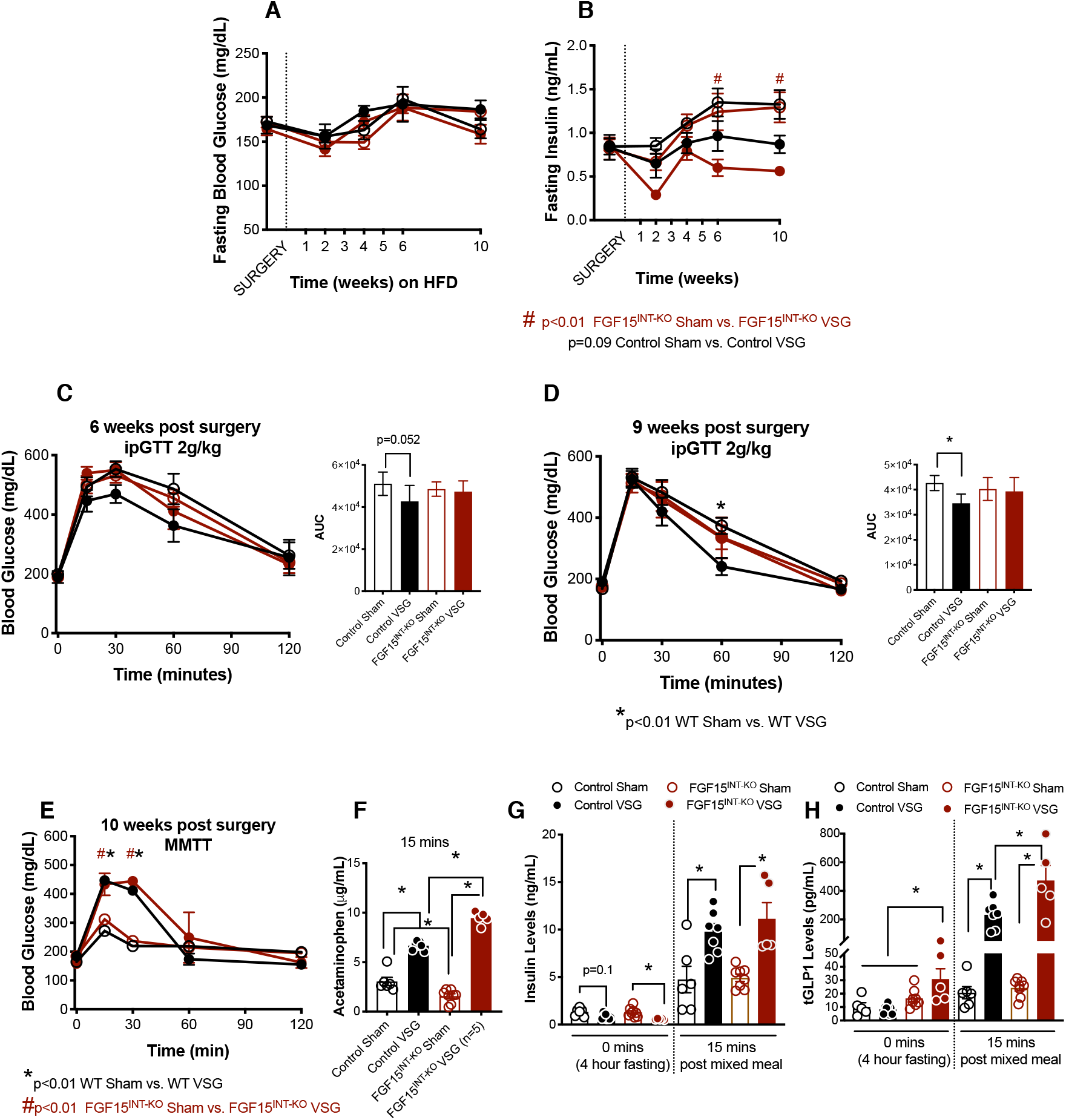
FGF15^INT-KO^ mice remain glucose intolerant despite significant body weight loss after VSG. **A.** Fasting (4 hours) blood glucose and **B**. Fasting (4 hours) insulin levels after surgery. **C** Intraperitoneal glucose tolerance test (ipGTT; 2g/kg) performed 6 weeks post-surgery and Area Under the Curve (AUC). **D** Intraperitoneal glucose tolerance test (ipGTT; 2g/kg) performed 9 weeks post-surgery and Area Under the Curve (AUC). **E.** Mixed meal tolerance test (MMTT) performed 10 weeks post-surgery. **F.** Gastric emptying rate measured by acetaminophen levels at 15 minutes post gavage. **G.** Insulin levels at baseline and 15 minutes post gavage. **H.** Total GLP-1 levels at baseline and 15 minutes post gavage. Animal number Control Sham (n=5-6), Control VSG (n=8), FGF15^INT-KO^ Sham (n=8), FGF15^INT-KO^ VSG (n=5). Data are shown as means ± S.E.M. *p<0.01 Control Sham vs. Control VSG; #p<0.01 FGF15^INT-KO^ VSG and FGF15^INT-KO^ Sham; ^p<0.01 Control VSG and FGF15^INT-KO^ VSG (2-Way ANOVA with Tukey’s post-test).

Next, we challenged mice with a mixed meal for the assessment of postprandial glucose excursion, insulin and GLP-1 response. Acetaminophen, which is rapidly absorbed once it leaves the stomach was added to the mixed meal to assess gastric emptying rate. FGF15^INT-KO^ Sham mice had a similar glucose excursion curve, but decreased gastric emptying rate compared to Control Sham mice (Figure 3E, F). Both VSG groups responded to the mixed meal with similar glucose excursion curves (Figure 3E), significantly higher than their Sham controls as a result of increased gastric emptying rate (Figure 3F). Despite similar glucose excursion, FGF15^INT-KO^ VSG mice showed significantly higher gastric emptying rate (indicated by the greater plasma acetaminophen levels) compared to Control VSG mice (Figure 3F). Basal and postprandial insulin levels were increased in both VSG groups compared to Sham controls (Figure 3G). Interestingly, basal and postprandial total GLP-1 levels were significantly higher in FGF15^INT-KO^ VSG mice compared to all other groups (Figure 3H).

### Loss of intestinal FGF15 results in aberrant hepatic lipid and glycogen metabolism following VSG

FGF15^INT-KO^ VSG had increased liver to body weight ratio (Figure 4A). Assessment of glycogen stores revealed that FGF15^INT-KO^ VSG have decreased liver glycogen and a trend toward higher skeletal muscle (tibialis anterior) glycogen content (Supplemental Figure 4A, B). Postprandial plasma alanine aminotransferase, ALT levels (predictor of liver injury) and plasma cholesterol levels were reduced in both VSG groups compared to Sham controls (Figure 4B, C), without significant changes in plasma free fatty acids or plasma triglycerides (Figure 4D, E).

**Figure 4.**
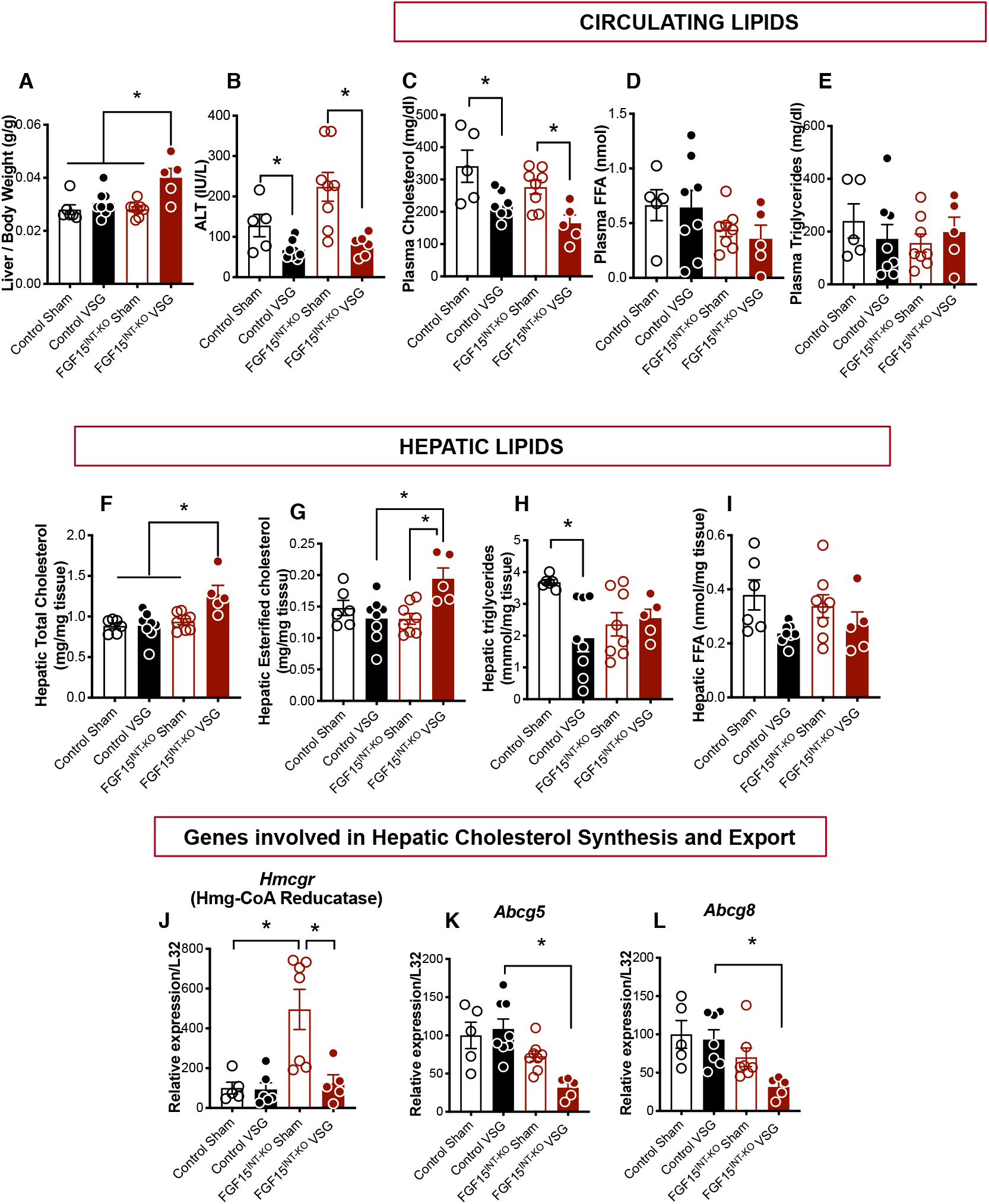
Loss of intestinal FGF15 results in aberrant hepatic lipid and glycogen metabolism following VSG. **A.** Liver weight normalized to body weight. **B.** Alanine aminotransferase (ALT) plasma levels. **C.** Plasma cholesterol **D.** Plasma free fatty acids (FFA) and **E.** Plasma triglycerides (postprandial, 12 weeks post-surgery). **F.** Total hepatic cholesterol. **G.** Hepatic esterified cholesterol. **H.** Hepatic triglycerides. **I**. Hepatic free fatty acids. Hepatic RNA expression of cholesterol synthesis gene **J.***Hmcgr* and cholesterol export genes **K.***Abcg5* and **L.***Abcg8.* Animal number Control Sham (n=5), Control VSG (n=8), FGF15^INT-KO^ Sham (n=8), FGF15^INT-KO^ VSG (n=5). Data are shown as means ± S.E.M. *p<0.05 (2-Way ANOVA with Tukey’s post-test).

Analysis of hepatic lipids revealed an increased cholesterol and esterified cholesterol levels in FGF15^INT-KO^ VSG mice (Figure 4F, G). Hepatic triacylglycerols were decreased in Control VSG compared to Control Sham mice, but remained at similar levels between FGF15^INT-KO^ Sham and FGF15^INT-KO^ VSG mice (Figure 4H). There was a trend toward decreased hepatic free fatty acids in both VSG groups compared to Sham controls (Figure 4I). Surprisingly, the cholesterol synthesis rate limiting gene, 3-hidroxy-3-methil-glutaryl-coenzyme A reductase, Hmg-CoA reductase (*Hmcgr*), was upregulated in FGF15^INT-KO^ Sham compared to Control Sham mice, but reduced after VSG in FGF15^INT-KO^ VSG mice (Figure 4J). Next, we measured the expression in hepatic cholesterol efflux pump-ATP-binding cassette, sub-family G, members 5 and 8 (*Abcg5* and *Abcg8*) and found that their expression was decreased in FGF15^INT-KO^ VSG mice (Figure 4K, L). These data suggest that despite decreased cholesterol synthesis, there is attenuated cholesterol export leading to elevated liver cholesterol content in FGF15^INT-KO^ VSG mice.

Hepatic expression of Farnesoid X receptor (*FXR*) was similar in Control and FGF15^INT-KO^ mice (Supplemental Figure 5A). We did see a trend of decreased expression of FXR in both VSG groups compared to their respective Sham groups (Supplemental Figure 5A). Fatty acid oxidation and lipid metabolism genes fatty acid synthase (*FAS*) and peroxisome proliferator-activated receptor alpha (*PPAR alpha*) were decreased in FGF15^INT-KO^ VSG mice compared to FGF15^INT-KO^ Sham controls (Supplemental Figure 5B, C). Although not significant, there was a trend of decreased expression of these genes in Control VSG compared to Control Sham mice (Supplemental Figure 5B, C). *PPAR alpha* target gene, Carnitine palmitoyltransferase 1 A (*Cpt1a*) essential for converting long-chain fatty acids into energy (lipogenesis), showed a trend toward decreased expression after VSG in FGF15^INT-KO^ but not Control mice (Supplemental Figure 5D). Interestingly, cluster of differentiation 36 (*CD36*), a cell surface protein that imports fatty acids, was increased in FGF15^INT-KO^ VSG compared to Control VSG mice (Supplemental Figure 5E).

### Intestinal FGF15 regulates enterohepatic bile acid metabolism following VSG

FGF15 is expressed in ileal enterocytes of the small intestine and released postprandially in response to bile acid absorption. Once released from the ileum, FGF15 enters the portal venous circulation and travels to the liver where FGF15 binds to its receptor FGFR4 and represses de novo bile acid synthesis (Inagaki et al., 2005). Consistent with this role to inhibit bile acid production, plasma and cecal content concentrations of bile acids are higher in mice lacking intestinal FGF15 (Figure 5A, B). VSG has been shown to alter both the concentration and composition of bile acids. Therefore, we assessed the role that FGF15 might play in the effect of VSG to alter bile acids. Lack of FGF15 in the FGF15^INT-KO^ Sham mice resulted in slight increase of circulating bile acid levels and a significant increase in the cecum content bile acid levels compared to Control Sham (Figure 5A, B). However, FGF15^INT-KO^ VSG mice had higher plasma bile acid levels, but normal cecum content bile acid levels when compared to Control VSG and FGF15^INT-KO^ Sham mice (Figure 5A, B). Not surprisingly, lack of intestinal FGF15 resulted in higher expression of bile acid synthesis gene *Cyp7a1* in FGF15^INT-KO^ Sham compared to Control Sham mice (Figure 5C). However, despite the fact that FGF15^INT-KO^ VSG mice lack intestinal FGF15 and have increased plasma bile acid levels, their expression of hepatic bile acid synthesis genes cholesterol 7a-hydroxylase *(Cyp7a1)*, sterol 12-alpha-hydroxylase *(Cyp8b1)* and sterol 27-hydroxylase *(Cyp27a1)* were decreased (Figure 5C, D, E).

**Figure 5.**
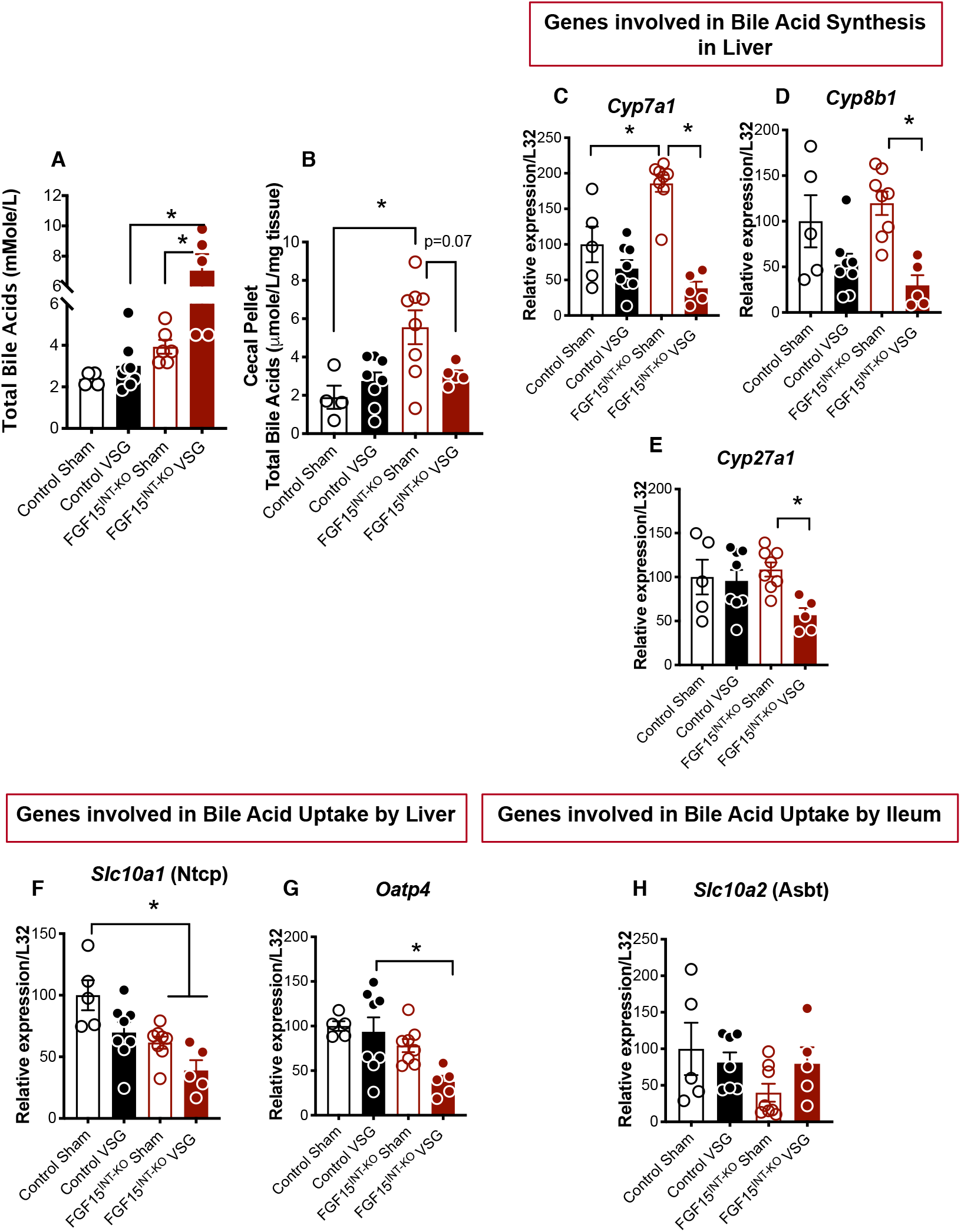
Intestinal FGF15 regulates enterohepatic bile acid metabolism following VSG. **A.** Total plasma bile acid levels. **B.** Cecal total bile acid levels. Hepatic RNA expression of bile acid synthesis genes **C.***Cyp7a1*, **D.***Cyp8b1* and **E.***Cyp27a1*. Hepatic RNA expression of bile acid uptake genes **F.***Slc10a1* (coding for Ntcp) and **G.***Oatp4*. **H.** Ileum RNA expression of bile acid uptake gene *Slc10a2* (coding for Asbt). Animal number Control Sham (n=4-5), Control VSG (n=8), FGF15^INT-KO^ Sham (n=8), FGF15^INT-KO^ VSG (n=5). Data are shown as means ± S.E.M. *p<0.05 (2-Way ANOVA with Tukey’s post-test).

Next, we measured the expression of hepatic and ileal bile acid uptake transporters. The expression of *Slc10a1* (coding for liver bile acid transporter LBAT/Ntcp) and *Oatp4* was decreased in FGF15^INT-KO^ VSG mice (Figure 5F, G). We did not see differences in the expression of bile acid transporter *Slc10a2* (coding for apical sodium-dependent bile acid transporter, Asbt) in the ileum (Figure 5H). These data suggest that bile acid uptake by liver is reduced in FGF15^INT-KO^ VSG mice, potentially contributing to the increased plasma bile acid levels.

### Intestinal FGF15 modulates microbiota in cecal content

Changes in gut microbiota composition are considered a potential contributor to the metabolic benefits of bariatric surgery (Basso et al., 2016; Liou et al., 2013; Sanmiguel et al., 2017). We investigated whether intestinal FGF15 modulates shifts in the microbial communities by VSG. To determine the effects of FGF15 on VSG-induced gut microbiota alteration, we performed 16S ribosomal RNA (rRNA) gene sequencing on cecal content samples collected 12 weeks after surgery (collected at time of sacrifice). Chao1 and Shannon index were used to estimate with-in sample richness and diversity, respectively. The former represents the total number of microbes present in one single sample, while the latter accounts for both richness and evenness of the microbes. We did not see significant changes in richness (Chao1 index) and diversity (Shannon index) (Figure 6A, B). However, there was a trend toward increased richness in both VSG groups compared to their respective Sham controls. Additionally, we observed a trend of decreased cecal diversity in FGF15^INT-KO^ Sham compared to Control Sham mice (Figure 6B).

**Figure 6.**
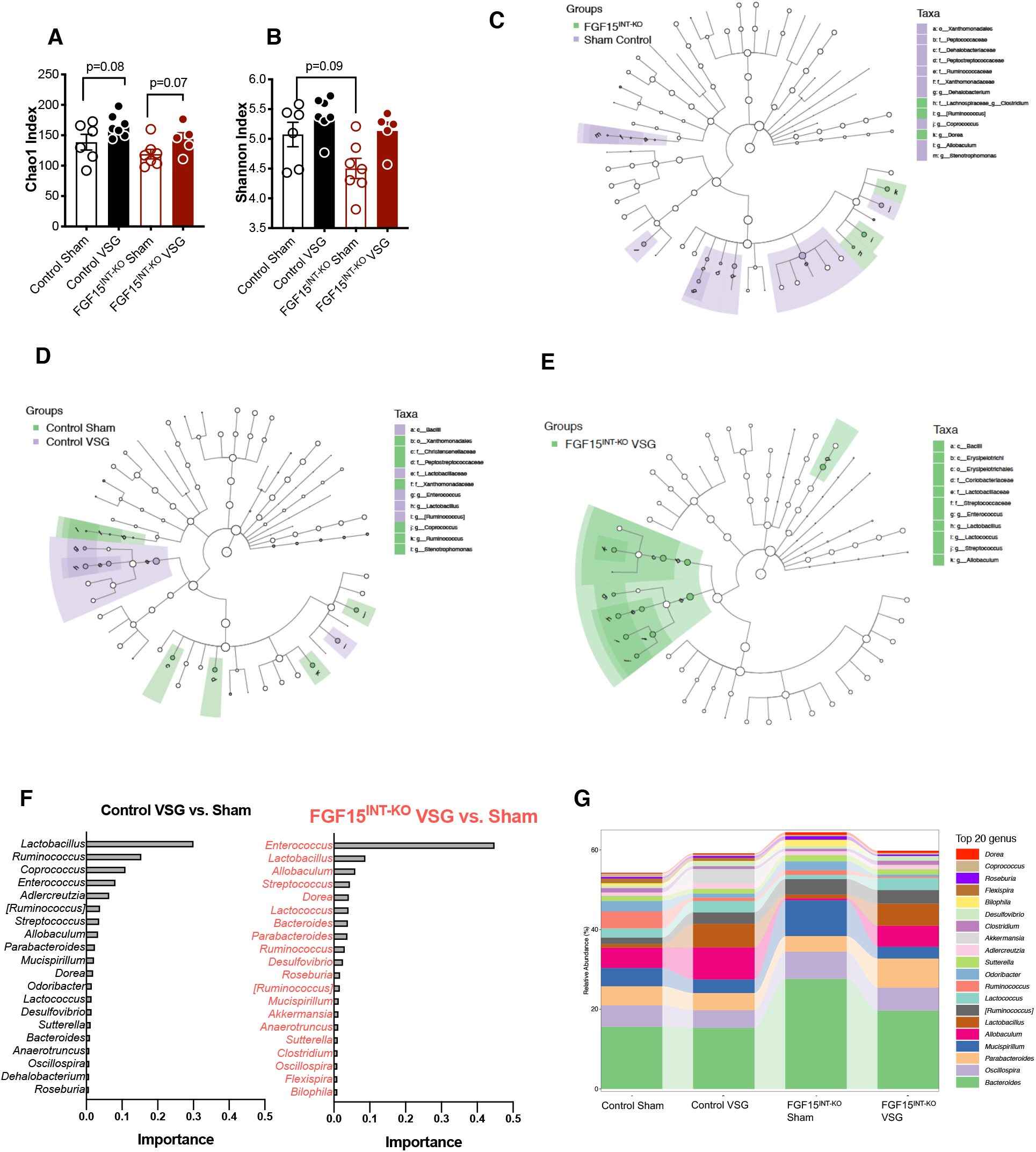
Intestinal FGF15 modulates microbiota in cecal content. **A.** Chao1 abundance and **B.** Shannon index diversity of the gut microbiota in the cecal contents. LEfSe analysis depicting nodes within the bacterial taxonomic hierarchy that are enriched in cecal microbiota from **C.** Control Sham versus FGF15^INT-KO^ Sham and **D.** Control Sham versus VSG and **E.** FGF15^INT-KO^ Sham versus VSG. Diagrams generated by LEfSe indicating differences at *phylum*, *order*, *class*, *family*, and *genus* levels between the two groups. **F.** Top ranked taxa at *genus* level identified by random forest analysis according to their ability to discriminate the microbiota of Sham and VSG mice per genotype in decreasing order of discriminatory importance. A comparison of the abundance of markers in VSG relative to Sham counterparts in each genotype. **G.** Differences in relative abundance of taxa at *genus* level. Animal number Control Sham (n=6), Control VSG (n=7), FGF15^INT-KO^ Sham (n=7), FGF15^INT-KO^ VSG (n=5). Data in A, B are shown as means ± S.E.M. *p<0.05 (2-Way ANOVA with Tukey’s post-test).

Linear discriminant analysis (LDA) effect size (LEfSe) analysis showed taxonomic differences in the microbiota composition of the cecal content between Control Sham and FGF15^INT-KO^ Sham groups (Figure 6C). *Firmicutes* and *Bacteroidetes* are the most abundant *phyla* in fecal microbiota. The relative proportion of *Firmicutes* and *Bacteroidetes* has been reported to be affected differently by obesity and high-fat feeding *Firmicutes* at the *phylum* level, *Erysipelotrichales* at *class* level, and *Allobaculum* and *Ruminococcus* at the *family* level showed a trend of decreased abundance in FGF15^INT-KO^ Sham compared to Control Sham (Supplemental Figure 6B, E, H, I). However, *Lachnospiraceae* at the *family* level and *Bacteroides* at the *genus* level were increased in FGF15^INT-KO^ Sham compared to Control Sham (Supplemental Figure 6F, G).

Linear discriminant analysis (LDA) effect size (LEfSe) analysis showed taxonomic differences in the microbiota composition of the cecal content between Sham and VSG in both genotypes groups (Figure 6D, E). To further define bacteria characteristic of enrichment of Sham and VSG in each genotype, we also performed a random forest model. *Lactobacillus* species have been suggested to confer beneficial effects in a broad spectrum (Crovesy et al., 2017; Wells and Mercenier, 2008). At the *genus* level, *Lactobacillus*, with an importance score of 0.301 determined by the decrease in the classification accuracy when it was ignored, was identified as the top-ranked discriminator for the Control VSG vs. Control Sham gut microbiota. On the other hand, *Enterococcus* genus was identified as the top discriminator for FGF15^INT-KO^ VSG vs. FGF15^INT-KO^ Sham gut microbiota, with *Lactobacillus* genus having an importance score of 0.088. (Figure 6F, G).

## Discussion

While obesity rates continue to rise, most treatment strategies such as dietary and lifestyle interventions are hampered by limited long-term efficacy. Weight loss surgeries, such as Roux-en-Y gastric bypass (RYGB) and Vertical Sleeve Gastrectomy (VSG), cause more weight loss and higher rates of diabetes resolution than other available therapies (Schauer et al., 2017). Increased circulating bile acids are one of the hallmarks of bariatric surgery, and our group and others have identified bile acid signaling through FXR as a molecular link mediating the beneficial metabolic effects of elevated bile acids following VSG (Albaugh et al., 2019; Ryan et al., 2014). Secretion of FGF15/19 is a key response to FXR activation. Consistent with human data that show increased circulating levels of FGF19 after bariatric surgery (DePaoli et al., 2019; Gomez-Ambrosi et al., 2017; Haluzikova et al., 2013; Mulla et al., 2019; Sachdev et al., 2016), VSG in mice upregulates FGF15 expression in the ileum (Figure 1). Here we demonstrate that FGF15 has important effects on metabolic, morphologic and bacteriologic responses to VSG.

To study the specific role of intestinal FGF15 in the response to VSG, we built a mouse model that allowed for specific deletion of FGF15 from the intestine in the adult animal (see Figure 1 and methods). Compared to control animals, these mice have similar body weight, body fat, food intake and glucose regulation when challenged on a high-fat diet. These data argue that intestinally derived FGF15 is not necessary for physiologic regulation of energy balance and glucose levels in diet-induced obese animals without bariatric surgery. However, when FGF15^INT-KO^ mice have VSG they actually lose considerably more weight than the control mice with surgery (Figure 1). Moreover, while we have consistently observed that mice given a VSG lose little or no lean mass, FGF15^INT-KO^ lose 25% of their lean tissue, much more than the control VSG mice (Figure 1). We have previously reported that wild type obese mice do not lose significant amount of lean body mass after VSG (Patel et al., 2018). Thus, while FGF15 does not mediate reductions in body fat after VSG, it is essential to protect from deleterious loss of lean mass with the rapid weight loss that occurs after VSG. Loss of muscle mass is a major concern in the setting of rapid weight loss interventions such as very-low calorie diets and bariatric surgery (Choksi et al., 2018; Gregory, 2017; Kenngott et al., 2019; Stein and Silverberg, 2014). In particular, loss in muscle mass can greatly inhibit the proper disposal of glucose; this may account for the failure of VSG to improve IP glucose tolerance in the FGF15^INT-KO^ mice (Figure 3).

A second component of lean tissue loss in the FGF15^INT-KO^ VSG mice was due to reduced bone mass (Figure 2). Bariatric procedures including VSG can result in bone loss beyond the normal response to reduced loading from weight loss, or calcium deficiency from diminished absorption (Gregory, 2017; Kim et al., 2015a; Li et al., 2019). In FGF15^INT-KO^ mice this effect of surgery is amplified and results in a diffuse pattern of osseous abnormality including both trabecular and cortical bone (Figure 2). In humans, reduction in bone density after bariatric surgery can predispose to fractures (Gregory, 2017). It remains to be seen whether this post-surgical metabolic bone disease can be accounted for by FGF15/19 signaling as suggested by our findings.

A recent study reported that exogenous FGF19 induces skeletal muscle hypertrophy and blocks muscle atrophy induced by glucocorticoid treatment, sarcopenia and obesity (Benoit et al., 2017). The mechanism behind the loss of muscle and bone mass in our FGF15^INT-KO^ VSG mice is not entirely clear and may include several scenarios. FGF21 levels were increased in FGF15^INT-KO^ mice (especially after VSG; Supplemental Figure 2). FGF21 levels are tighly regulated to nutrional status, and it has been well established that FGF21 levels increase in fasting and after ketogenic and low-protein diets (Badman et al., 2007; Inagaki et al., 2007; Pezeshki et al., 2016). The lack of FGF15, which is a postprandial hormone, could be altering the gut-liver communication and leading to the liver sensing a “fasting state” and increasing FGF21 levels. Alternatively, other tissues, including muscle, could be responsible for the increased FGF21 levels in FGF15^INT-KO^ VSG mice (Tezze et al., 2019). This is consistent with the observation that elevated plasma FGF21 levels have been linked to decreased muscle mass (Oost et al., 2019) and reduced bone mineral density and BMAT (Bornstein et al., 2014; Fazeli et al., 2015; Wei et al., 2012). Therefore, increased FGF21 levels could be responsible for the muscle and bone loss in FGF15^INT-KO^ VSG mice.

The critical point, however, is that increased levels of intestinal-derived FGF15 protect from potential deleterious effects of rapid weight loss including loss of muscle and bone. These data are important in the context of patients undergoing weight-loss surgery. Current strategies to mitigate these effects include adjustments in diet, weight resistance training as well as calcium and vitamin D supplementation before and after surgery (Krez and Stein, 2020). Our data suggest that bariatric surgery patients with low FGF19 levels may be at a higher risk for bone and skeletal muscle loss. More research is warranted to determine if FGF19 can act as a biomarker to identify patients that are at high risk for excessive muscle and bone loss following bariatric surgery. Understanding this relationship would allow physicians to optimize treatment strategies for at risk patients.

We did not see a difference in glucose excursion between Control Sham and FGF15^INT-KO^ Sham mice, suggesting that intestinal-derived FGF15 does not play a role in peripheral glucose tolerance, at least under high-fat diet conditions (Figure 3). As expected, we saw an improvement in glucose tolerance in Control VSG compared to Control Sham mice after intraperitoneal glucose challenge (ipGTT; Figure 3). To our surprise, there was no difference in glucose tolerance between FGF15^INT-KO^ VSG and FGF15^INT-KO^ Sham mice, despite the large body weight loss in FGF15^INT-KO^ after VSG (Figure 3). Loss in muscle mass as seen in FGF15^INT-KO^ VSG can greatly inhibit the proper disposal of glucose, leading to glucose intolerance. Recent studies reported that patients who experience post-bariatric postprandial hypoglycemia have increased postprandial levels of FGF19, linking the levels of FGF15/19 to glucose regulation after weight-loss surgery (Mulla et al., 2019). Despite glucose intolerance, FGF15^INT-KO^ VSG mice had lower hepatic glycogen content compared to FGF15^INT-KO^ Sham mice (Supplemental Figure 5). This suggests that the liver is sensing a relative state of starvation. Interestingly, we also observed a trend of increased glycogen content in skeletal muscle of FGF15^INT-KO^ VSG mice (Supplemental Figure 4). This points toward the physiological importance of intestinal FGF15 on glycogen metabolism in liver and muscle after VSG. As mentioned, the literature has been clear on the positive role of FGF15 on hepatic glycogen content (Kir et al., 2011). However, it is unclear whether lack of FGF15 in VSG mice directly affects skeletal muscle glycogen content or if our observations are secondary to the decreased lean muscle mass in these mice.

Reduced expression of FGF15 in mice leads to increased gastrointestinal motility, increased plasma levels of bile acids and increased luminal water content, similar to human bile acid diarrhea (Lee et al., 2018). However, gastric emptying was lower in FGF15^INT-KO^ Sham compared to Control Sham mice (Figure 3). These differences in gastric emptying rates change after VSG. FGF15^INT-KO^ VSG had increased gastric emptying compared to Control VSG mice (Figure 3). Consistent with previous reports, we also saw an increased total GLP-1 postprandial response in Control VSG mice after a mixed meal. The increase in total GLP-1 after VSG has been attributed to the rapid entry of ingested glucose and nutrients into the small intestine (Jorgensen et al., 2015). However, the GLP-1 (basal and postprandial) in FGF15^INT-KO^ VSG mice were much higher than Control VSG (Figure 3). This points to a more complicated regulation of GLP-1 secretion beyond nutrient presentation. Strong evidence links bile acid activation of TGR5 to increased GLP-1 secretion (Katsuma et al., 2005; Potthoff et al., 2013; Thomas et al., 2009). Consequently, elevated levels of bile acids in FGF15^INT-KO^ mice given VSG may act to increase TGR5 signaling and drive increased GLP-1 secretion. Interestingly, despite the elevated GLP-1 levels, FGF15^INT-KO^ mice given VSG do not have improved glucose tolerance pointing to a potentially more central role of FGF15 as compared to GLP-1.

Another weight-independent effect of bariatric surgery is the changes in enterohepatic bile acid circulation resulting in increased plasma bile levels as well as altered bile acid composition in rodents (Kohli et al., 2010; Myronovych et al., 2014) and humans after bariatric surgery (Patti et al., 2009; Pournaras et al., 2012). FGF15/19 is released postprandially in response to bile acid absorption (Potthoff et al., 2011). Once released from the ileum, FGF15/19 enters the portal venous circulation and travels to the liver where FGF15/19 binds to its receptor FGFR4 and represses de novo bile acid synthesis through suppression of the rate limiting enzyme cholesterol 7a-hydroxylase (*Cyp7a1*). Therefore, bile acids and FGF15 inhibit further bile acid synthesis and facilitate communication between the liver and small intestine. As predicted, lack of FGF15 resulted in increased plasma bile acids and cecal bile acid content in FGF15^INT-KO^ Sham mice (Figure 5). After VSG, plasma bile acids were further increased in FGF15^INT-KO^ VSG mice, but cecal bile acids levels were suppressed (Figure 5). This suggests that VSG directly alters the compartmental-specific bile acid pool independently of FGF15.

Patients with NAFLD have increased hepatic *Cyp7a1* levels (DePaoli et al., 2019). Exogenous administration of FGF19 and the analog NGM282 did not correct hyperglycemia in diabetic patients, but caused a rapid and sustained reduction in hepatic *Cyp7a1* levels and liver fat content in NAFLD patients (DePaoli et al., 2019). Consistent with these observations, we saw increased hepatic expression of *Cyp7a1* in FGF15^INT-KO^ Sham mice compared to Control Sham (Figure 5). However, hepatic *Cyp7a1* was significantly downregulated in FGF15^INT-KO^ VSG (Figure 5). We speculate that the drastic increase in plasma bile acid levels in FGF15^INT-KO^ VSG mice downregulates hepatic *Cyp7a1* expression as a negative feedback, despite lack of circulating FGF15 (Agellon and Cheema, 1997; Baker et al., 2000). We also speculate that these pathways are independent of FXR. A recent study showed that FGF19 analog modulates bile acid homeostasis even when administered to FXR knock out mice (Gadaleta et al., 2020). Bile acids induce *PPAR alpha* transcription via induction of FXR and we did not see difference in the RNA levels of either in FGF15^INT-KO^ compared to control mice (Supplemental Figure 5) (Pineda Torra et al., 2003). We also did not see FGF15-dependent regulation of *Cyp8b1* expression, as it has been suggested before that its expression is regulated by FXR/SHP signaling and not FXR/FGF15/FGFR4 pathway (Figure 5)(Kim et al., 2007).

Future studies will need to dissect the signaling pathways responsible for the decreased *Cyp7a1* expression in FGF15^INT-KO^ VSG mice. Understanding the pathways that suppress *Cyp7a1* would also be useful for cancer treatments that aim to block FGFR4. FGF19 signaling has been implicated in the development of hepatocellular carcinoma, making FGFR4 antagonists attractive candidates for treating this disease (Schadt et al., 2018). Blockade of FGFR4 signaling inhibits the FGF19-induced bile acid brake. The resulting elevation in hydrophobic bile acids that act as detergents and disrupt cell membranes may lead to liver damage (Perez and Briz, 2009). Although we see significantly elevated levels of bile acids in the FGF15^INT-KO^ VSG compared to Control VSG mice, we do not observe signs of liver damage as noted by normal circulating levels of ALT in FGF15^INT-KO^ VSG mice (Figure 4). The VSG-induced reduction of *Cyp7a1* may prevent further increase in bile acid levels and subsequent bile acid toxicity that is noted with reduced FGF4R signaling. Additionally, VSG and lack of FGF15 (FGF15^INT-KO^ Sham and VSG mice; Figure 5) showed reduced levels of hepatic *Slc10a1* (encoding for Ntcp), which has been recently shown as a potential target for the treatment of obesity and fatty liver disease (Donkers et al., 2020).

These data show that intestinal FGF15 is necessary for the reduction in hepatic cholesterol content after VSG. FGF15^INT-KO^ VSG mice had increased liver weight (normalized to body weight), despite decreased hepatic glycogen content (Figure 4 and Supplemental Figure 5). Although plasma cholesterol levels were decreased in both groups after VSG, FGF15^INT-KO^ VSG mice had increased hepatic total and esterified cholesterol levels (Figure 4). The cholesterol synthesis rate limiting gene, Hmg-CoA reductase (*Hmcgr*), was upregulated in FGF15^INT-KO^ Sham compared to Control Sham mice, but surprisingly reduced after VSG in FGF15^INT-KO^ VSG mice (Figure 4). Hepatic cholesterol efflux pump-ATP-binding cassette, sub-family G, members 5 and 8 (*Abcg5* and *Abcg8*) expression was decreased in FGF15^INT-KO^ VSG mice (Figure 4). These data suggest that despite decreased cholesterol synthesis there is attenuated cholesterol export leading to elevated liver cholesterol content in FGF15^INT-KO^ VSG mice. Although, hepatic expression of lipogenic genes *FAS*, *PPAR alpha* and lipolytic gene *Cpt1a* were lower in FGF15^INT-KO^ VSG compared to FGF15^INT-KO^ Sham mice, *CD36* was increased in FGF15^INT-KO^ VSG versus Control VSG liver (Supplemental Figure 5). Overexpression of the lipogenic gene *CD36* is associated with lipoprotein uptake and increased steatosis in the liver of patients with NAFLD (Bechmann et al., 2010). Exogenous FGF19 suppresses hepatic *CD36* increase, contributing to its role in lipid-mediated cellular stress and liver injury (Alvarez-Sola et al., 2017).

Clinical and mouse studies have reported increased plasma FGF21 levels in patients and animal models of NAFLD (Dushay et al., 2010; Li et al., 2010; Zhang et al., 2008). Consistent with these data, we believe that plasma FGF21 is increased in FGF15^INT-KO^ mice as compensatory mechanism to attenuate liver injury and hepatic lipid accumulation (Kim et al., 2015b). Importantly, we did not observe increased hepatic cholesterol levels in FGF15^INT-KO^ Sham mice compared to Control Sham (despite increased hepatic expression of *Hmcgr*). It is possible that the hepatic cholesterol in these mice is directed towards their increased plasma bile acid levels. It is also possible that increased FGF21 levels in FGF15^INT-KO^ Sham were sufficient to counteract the hepatic lipid accumulation and liver damage. Our hypothesis is that the weight loss surgery dependent increase in plasma/intestinal FGF15 and plasma bile acids has a role in mediating the surgery’s potent effect on enterohepatic metabolism, specifically the regulation of bile acids and hepatic lipid synthesis.

The gut microbiota is an important regulator of bile acid metabolism, regulating the synthesis of bile acids and production of secondary bile acids (Swann et al., 2011). The primary bile acids (chenodeoxycholic acid and cholic acid) are actively reabsorbed in the ileum, but those that escape reabsorption are deconjugated to deoxycholic acid and lithocholic acid by colonic bacteria and reabsorbed through the portal system. The colonic bacteria involved in the deconjugaton of bile acids are mostly *Bacteroides* species, which studies have found to be decreased in bariatric surgery patients (Damms-Machado et al., 2015; Dewulf et al., 2013) and rodents (Ryan et al., 2014) and this change is correlated with decreased adiposity and improved glucose control. FGF15^INT-KO^ Sham mice had increased abundance of *Bacteroides* in cecal microbiota, which was reduced after VSG (Figure 6 and Supplemental Figure 6). The increased cecal bile acids observed in FGF15^INT-KO^ Sham were also reduced after VSG (Figure 5). We hypothesize that the reduction in *Bacteroides* could contribute to the reduction in cecal bile acids in FGF15^INT-KO^ after VSG, showing that VSG alters the gut microbiome in the absence of FGF15. We also saw a trend of increased *Bacteroidetes* and decreased *Firmicutes* at the phyla level in FGF15^INT-KO^ Sham mice, similar to what has been reported in FXR knock out mice (Supplemental Figure 6)(Parseus et al., 2017). Additionally, *Lachnospiraceae* at the family level was increased in FGF15^INT-KO^ Sham mice (Supplemental Figure 6). The changes in these taxa are consistent with resistance to diet-induced obesity (Lecomte et al., 2015). That is surprising since both Sham groups were nearly identical in body weight and glucose homeostasis. We also investigated how intestinal FGF15 modulates shifts in the microbial communities by VSG. *Lactobacillus* species have been suggested to confer beneficial effects in a broad spectrum of situations (Crovesy et al., 2017; Wells and Mercenier, 2008). Random forest test analysis showed that *Lactobacillus* was the main driver of the difference between Control Sham and VSG at the order level, where it was much lower in importance in FGF15^INT-KO^ groups (Figure 6D). *Enterococcus* was the main driver of difference between the FGF15^INT-KO^ Sham and VSG groups at the *genus* level (Figure 6D). Increase in *Enterococcus* species in the gut has been linked to decreased adiposity as a result of increased energy expenditure (Quan et al., 2019). A recent study showed that a probiotic cocktail of *Lactobacillus* and *Enterococcus* prevented diet-induced inflammation and leaky gut by increasing bile acid hydrolase activity (Ahmadi et al., 2020). Overall, this data supports the independent roles of FGF15 and VSG in the modulation of gut microbiota and future studies will dissect these roles on metabolism and energy expenditure.

In conclusion, our findings point to an important role for FGF15 in the regulation of multiple metabolic parameters following VSG. Intestinal-derived FGF15 increases in mouse models of VSG and is necessary for the improvement in blood glucose regulation, to preserve muscle mass and bone mass, and for the decrease in hepatic cholesterol after VSG-induced weight loss. Our data also shows that the VSG-induced increase in intestinal FGF15 has a specific role in mediating the surgery’s potent effect on enterohepatic metabolism, specifically the regulation between bile acids, hepatic lipid synthesis and gut microbiome.

## RESOURCE AVAILABILITY

### Lead Contact

Further information and requests for resources should be directed towards Dr. Randy J. Seeley.

### Materials Availability

Reagents and genetically modified mice developed in the context of this manuscript will be shared with investigators who request them in accordance with institutional guidelines using a simple Material Transfer Agreement. Mouse models prior to publication will be available to the requesting investigator as a collaboration (this would be determined on a case-by-case basis). Animals that have been published will be made available to the scientific community at-large upon request or through deposition into central repositories.

### Data and code availability

This study did not generate unique datasets or code.

## EXPERIMENTAL MODEL AND SUBJECT DETAILS

### Study Approval

All protocols were approved by the University of Michigan (Ann Arbor, MI) Animal Care and Use Committees and were in accordance to NIH guidelines.

### Generation of FGF15 flox mouse

Gene sequence for the mouse FGF15 gene (4.4k base pair double strands DNA containing fgf15 exon) was downloaded from genebank. Guide RNAs were designed against the mouse sequence containing the region of interest for targeting (introns flanking FGF15 exon 2) using prediction algorithms available through http://crispor.tefor.net/. Guides were selected based on the specificity score and predicted efficiency (Mor.Mateos). Selected single guide RNA (sgRNA) sequences were subcloned into plasmid pX330 and subsequently confirmed by sequencing. Targeting DNA oligonucleotides including the 34 base pairs loxP site and 81-82 nucleotides of flanking sequence on either side were generated (IDT Technologies). DNA oligonucleotides were designed to disrupt the sgRNA and PAM sequences so that Cas9 would not be able to cleave the inserted sequence after incorporation into the genome. To test the ability of the sgRNAs to cut chromosomal DNA appropriately, each sgRNA was injected into fertilized eggs by the University of Michigan Transgenic Animal Core; zygotes were then allowed to develop into blastocysts in culture. PCR amplification of the target region followed by sequencing was used to confirm Cas9 dependent DNA cutting in vivo. Cas9/sgRNA/oligo donor were then injected into 300 fertilized mouse eggs (C57BL/6x SJL) and transferred to pseudo-pregnant recipients for gestation. After delivery of potential founders, tail DNA was isolated and screened by PCR across the region of interest to identify genomic manipulations. As founders are often mosaic for allelic changes, PCR products of genomic DNA were subcloned into topo vector and sequenced to characterize the genetic modification in the mice.

### Animals and Diet

The FGF15 flox/flox mice, were built using CRISPR-Cas9 technology with LoxP sites flanking exon 2 of the FGF15 gene (described above). We bred these mice to VilCreERT2 mice and administered tamoxifen (intraperitoneal, 3 doses/150 mg/kg) to VilCreERT2; Fgf15^flox/flox^ and controls (VilCreERT2 and Fgf15 flox/flox). We validated exon 2 excision within latter jejunal and ileal mucosa, where FGF15 is most highly expressed (Supplemental Figure 1). Male mice were single-housed under a 12-hour light/dark cycle with ad libitum access to water and food. A week after tamoxifen administration, FGF15^INT-KO^ and Control male mice (all littermates) were placed on 60% HFD from Research Diets, Inc. (New Jersey, US; Catalog D12492) for 8 weeks. Mice underwent VSG or Sham surgery (described below) and returned to 60% HFD four days after surgery until end of study.

Animals were euthanized 12 weeks post-surgery. One Control Sham mouse accidently died during NMR measurements on the day before necropsy. The post-necropsy data on metabolites and tissue gene expression for this Control Sham mouse was excluded. The rest of the mice were fasted overnight, administered oral mixed meal (volume 200 μl Ensure Plus spiked with a 40-mg dextrose) and sacrificed 90 minutes later. Plasma and tissues were collected and frozen immediately. All animals were euthanized using CO_2._

### Vertical Sleeve Gastrectomy (VSG) in mice and rats

Mice were maintained on a 60% HFD for 8 weeks prior to undergoing Sham or VSG surgery, as described previously (Evers et al., 2019; Kim et al., 2019; Patel et al., 2018). Mice were fasted overnight prior to day of surgery. Animals were anesthetized using isoflurane, and a small laparotomy incision was made in the abdominal wall. The lateral 80% of the stomach along the greater curvature was excised in VSG animals by using an ETS 35-mm staple gun (Ethicon Endo-Surgery). The Sham surgery was performed by the application of gentle pressure on the stomach with blunt forceps for 15 seconds. All mice received one dose of Buprinex (0.1 mg/kg) and Meloxicam (0.5 mg/kg) immediately after surgery. All mice received Meloxicam (0.5 mg/kg) for 3 days after surgery and Enrofloxacin (40 mg/kg) for 5 days after surgery. Animals were placed on DietGel Boost (ClearH_2_O; Postland ME) for 3 days after surgery. They were placed back on pre-operative solid diet (60% HFD) on day 4 post-surgery. Body weight and food intake as well as overall health were monitored daily for the first 7 days after surgery and once weekly until end of the studies.

### Metabolic Studies

Body weight was monitored monthly for 9 weeks prior and 12 weeks after Sham/VSG surgery. Intraperitoneal glucose tolerance test (IPGTT) was performed by intraperitoneal (IP) injection of 50% dextrose (2g/kg) in 4-hour fasted male mice. Mixed-meal tolerance test (MMTT) was performed via an oral gavage of liquid meal (volume 200 μl Ensure Plus spiked with a 40-mg dextrose) in 4-hour fasted male mice. Blood was obtained from the tail vein and blood glucose was measured with Accu-Chek blood glucose meter (Accu-Chek Aviva Plus, Roche Diagnostics). Ten weeks after surgery, the rate of gastric emptying was assessed as previously described (Chambers et al., 2014). Briefly, after 4-hour fast, we delivered liquid mixed meal orally (volume 200 μl Ensure Plus spiked with a 40-mg dextrose and 4-mg acetaminophen, Sigma-Aldrich). Blood was collected from the tail vein at baseline and 15 min after gavage in EDTA-coated microtubes. Plasma acetaminophen levels were measured using spectrophotometry (Sekisui Diagnostics).

## METHOD DETAILS

### ELISA and Metabolite Assays

Insulin (Crystal Chem) and total GLP-1 (MesoScale Discovery) were measured during experiments shown in Figure 3. Postprandial plasma obtained at termination of studies (see above for details) was used to measure IGF-1 (R&D Systems), Activin-A (R&D Systems), Myostatin (GDF8) (R&D Systems), FGF21 (R&D Systems); total bile acids in plasma and cecal samples was measured with Total Bile Assay (NBT Method) assay (GenWay Biotech Inc; San Diego, CA). Glycogen content was measured in liver and muscle (tibialis anterior) samples with Glycogen Assay kit (Sigma-Aldrich). All sampled blood was collected via tail vein in EDTA-coated tubes. All assays were performed according to the manufacturer’s instructions.

### Lipid Measurements

Liver lipids were extracted with Lipid Extraction Kit Chloroform Free (Abcam). Total and cholesterol ester (Millipore/Sigma-Aldrich), triglycerides (Abcam), free fatty acids (Pointe Scientific) were measured using the extracted liver lipids. Postprandial plasma obtained at termination of studies (see above for details) was used to measure total cholesterol (Pointe Scientific), triglycerides (Pointe Scientific), free fatty acids (Pointe Scientific), ALT (Pointe Scientific). All assays were performed according to the manufacturer’s instructions.

### Bone Parameters

Tissues were fixed in 10% neutral-buffered formalin for 24 hours and kept in Sorenson’s buffer (pH7.4) thereafter. Tibiae were placed in a 19-mm diameter specimen holder and scanned over the entire length of the tibiae using a microcomputed tomography (μCT) system (μCT100 Scanco Medical). Scan settings were as follows: voxel size 12 μm, 70 kVp, 114 μA, 0.5 mm AL filter, and integration time 500 ms. Density measurements were calibrated to the manufacturer’s hydroxyapatite phantom. Analysis was performed using the manufacturer’s evaluation software and a threshold of 180 for trabecular bone and 280 for cortical bone. Tibiae used for μCT scanning were decalcified in 14% EDTA for 3 weeks. Paraffin-embedded tissue sections were processed and stained with H&E.

Bone Marrow Adipose Tissue Quantification by Osmium Tetroxide Staining and μCT: Mouse tibiae were decalcified in 14% EDTA for 2-3 weeks, and then put into 1% osmium tetroxide solution (diluted by Sorenson’s buffer pH7.4) for 48 hours. Osmium tetroxide-stained bones were scanned by the same program as described above. A threshold of 400 Gy was used for BMAT quantification. The volume of BMAT was normalized by the total volume (TV) of bone and shown as percentage (%).

### Muscle Fiber Area and Ileal Crypt Depth/Villi Height Analysis

Soleus muscle and ileal section were dissected and fixed in 10% neutral buffered formalin overnight. Tissue was embedded in paraffin and sectioned onto slides, and stained for H&E following standard protocol. Photos, and analysis of muscle fiber area (in 100-250 muscle fibers), and ileal crypt depth and ileal villi height (in 25 villi and crypts) were acquired using Olympus IX73 fluorescence microscopy system (Olympus). Villus height was measured from the crypt-villus junction to the tip of the villus and crypt depth was measured from the base of the crypt to the crypt-villus junction. Images were analyzed using Olympus cellSens imaging software (Olympus).

### Grip Strength

Grip strength was measured with Columbus Instruments Grip Strength Meter, which assesses neuromuscular function by sensing the peak amount of force an animal applies in grasping specially designed pull bar assemblies. Metering was performed with precision force gauges in such a manner as to retain the peak force applied on a digital display. The dual sensor model was employed by first allowing the animal to grasp the forelimb pull bar assembly. The animal was then drawn along a straight line leading away from the sensor. The animal released at some point and the maximum force attained is stored on the display. Each animal was tested five times and the average force reported in the data. Grip strength test was performed by the University of Michigan Physiology Phenotyping Core.

### Absorbed Energy Content

Fecal energy was assessed by University of Michigan Animal Phenotyping core using Bomb Calorimeter, Parr 6200 and 1108P oxygen bomb, as previously described (Asai et al., 2013). Mice were single-housed in clean housing cages for one week. Food weight was determined for the same time period. All fecal samples are collected using forceps and weighed to determine “wet weight”. Prior to processing, fecal samples are dried overnight at 50°C. Samples are then removed from the oven one at a time and weighed. Sample weights are recorded and fecal samples are then ground up individually. Samples are ground with mortar and pestle and carefully scooped into a dry tube. All instruments are washed with sparkleen and 10% bleach and completely dried using chem-wipes or paper towel between samples. New weighboats are used for each sample and weighed on the same scale as the pre-dried weights.

### Quantitative Real-Time PCR

RNA was extracted from tissue samples using RNeasy isolation kit (Qiagen). cDNA was synthesized by reverse transcription from mRNA using the iScript cDNA Synthesis Kit (Bio-Rad). Gene expression was performed by quantitative real time RT-PCR using Taqman gene expression assay was performed using StepOnePlus detection system (Applied Biosystems) with a standard protocol. Relative abundance for each transcript was calculated by a standard curve of cycle thresholds and normalized to RL32.

#### Intestinal Biometry

Following euthanasia, the entire gastrointestinal tract from the stomach to the rectum was removed, cleaned of mesenteric fat and gut weight and length determined. Small and large intestine/colon length was measured on a horizontal ruler after flushing with PBS. The entire small and large intestine/colon were then blotted to remove PBS before being weighed.

#### Analysis of 16S rRNA Gene Sequences

##### 16S rRNA Sequencing

Cecal contents were added to individual Bead plate provided by the Microbiome Core at the University of Michigan. DNA was isolated using Qiagen MagAttract PowerMicrobiome kit DNA/RNA kit (Qiagen, catalog no. 27500-4-EP) on the EpMotion 5075 (Eppendorf) liquid handler. Extracted DNA was then used to generate 16S rRNA libraries for community analysis. The DNA libraries were prepared by the Microbiome Core as described previously (Seekatz et al., 2015). Briefly, DNA was PCR amplified using a set of barcoded dual-index primers specific to the V4 region of the 16S rRNA gene (Kozich et al., 2013). PCR reactions are composed of 5 μL of 4 μM equimolar primer set, 0.15 μL of AccuPrime Taq DNA High Fidelity Polymerase, 2 μL of 10x AccuPrime PCR Buffer II (Thermo Fisher Scientific, catalog no.12346094), 11.85 μL of PCR-grade water, and 1 μL of DNA template. The PCR conditions used consisted of 2 min at 95°C, followed by 30 cycles of 95°C for 20 seconds, 55°C for 15 seconds, and 72°C for 5 minutes, followed by 72°C for 10 min. Each PCR reaction is normalized using the SequalPrep Normalization Plate Kit (Thermo Fisher Scientific, catalog no. A1051001). The normalized reactions are pooled and quantified using the Kapa Biosystems Library qPCR MasterMix (ROX Low) Quantification kit for Illumina platforms (catalog no. KK4873). The Agilent Bioanalyzer is used to confirm the size of the amplicon library (~399 bp) using a high-sensitive DNA analysis kit (catalog no. 5067-4626). Pooled amplicon library is then sequenced on the Illumina MiSeq platform using the 500 cycle MiSeq V2 Reagent kit (catalog no. MS-102-2003) according to the manufacturer’s instructions with modifications of the primer set with custom read 1/read 2 and index primers added to the reagent cartridge. The “Preparing Libraries for Sequencing on the MiSeq” (part 15039740, Rev. D) protocol was used to prepare libraries with a final load concentration of 5.5 pM, spiked with 15% PhiX to create diversity within the run. FASTQ files are generated when the 2 x 250 bp sequencing completes.

##### Analysis

Following sequencing, microbiome bioinformatics were run using QIIME 2 2020.2 (Bolyen et al., 2019). Briefly, non-singleton amplicon sequence variants (ASVs, 100% operational taxonomic units (OTUs)) were generated from raw sequences after trimming with the cutadapt plugin denoising with the dada2 plugin. One Control VSG and one FGF15^INT-KO^ Sham samples were excluded because of low OTUs. Taxonomy was then assigned to ASVs using the classify-sklearn alignment algorithm (Bokulich et al. 2018) against the Greengenes database (Release 13.8) of 99% OTUs reference sequences (McDonald et al. 2012). Alpha diversity metrics including Chao1 and Shannon, which estimate with-in sample richness and diversity respectively, were calculated using the diversity plugin. Chao1 index represents the number of ASVs present in one single sample, while Shannon index accounts for both abundance and evenness of ASVs present. Beta diversity metrics including weighted and unweighted UniFrac distance matrix (Lozupone et al., 2007), which estimate between-sample dissimilarity, were scaled and visualized through principle coordinates analysis (PCoA), and further used to determine the significance of the clustering between groups via permutational multivariate analysis of variance (PERMANOVA). Linear discriminant analysis (LDA) effect size (LEfSe) with default parameters (Segata et al., 2011) and Random Forest Classifier with 10-fold cross-validations (Breiman, 2001) were computed to identify significantly different microbes in abundance between groups at different taxonomic levels.

## QUANTIFICATION AND STATISTICAL ANALYSIS

The statistical analysis for comparisons between 2 groups was performed by unpaired (2-tailed) Student’s t test. Two-way ANOVA with post hoc Tukey’s multiple comparisons test was used for comparisons among 4 groups. P values <0.05 were considered significant (GraphPad Prism 8.2.0). Microbiome analysis is described above.

## Acknowledgements

The authors thank the surgeons for conducting mouse VSG (Alfor Lewis, Andriy Myronovych, Mouhamadoul Toure) and Kelli Rule, Jack Magrisso and Stace Kernodle for the technical assistance. We thank Dr. Dan Michele for discussion of the data (University of Michigan). We thank University of Michigan Animal Phenotyping Core (1U2CDK110678-01), University of Michigan Physiology Phenotyping Core (P30-AR069620), Adipose Tissue Core of the MNORC (P30 DK089503), Michigan Integrative Musculoskeletal Health Core Center (P30 AR069620) and University of Michigan In-Vivo Animal Core (IVAC) at University of Michigan, Ann Arbor. This work was supported by Novo Nordisk (RJS), NIH 5T32DK108740 (NB), 5T32DK071212-12 (NB), DK020572 (MDRC), DK089503 (MNORC), DK107282 (DAS), DK121995 (DAS), RO1 DK62876 (OAM), R24 DK092759 (OAM), China Scholarship Council grant (CSC#201606100218) to YS, and American Diabetes Association (1-18-PDF-087) to ZL.

## Author Contributions

NB and RJS conceived, designed, analyzed results and wrote the manuscript. NB, JS, YS, RGA, SC, SGV, ZL, KG performed the experiments and analyzed results. KMH, OAM and DAS helped with data interpretation and discussion. All authors edited manuscript. RJS provided final approval of the submitted manuscript.

