## Supplemental Tables and Figures for "Intestinal-derived FGF15 preserves muscle and bone mass following sleeve gastrectomy"

### Supplemental Figure 1

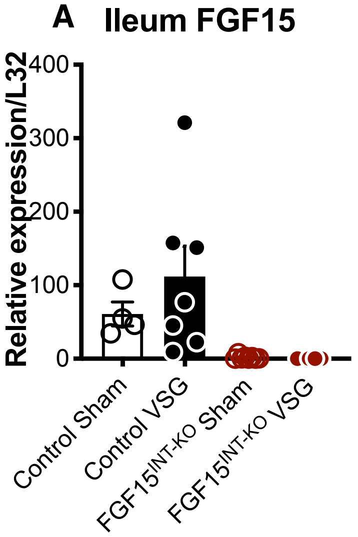


**Supplemental Figure 1. The expression of ileal FGF15 expression is ablated in FGF15$^{INT\text{-}KO}$ mice. A.** RNA expression of FGF15 in the ileum. Animal number WT Sham (n=5), WT VSG (n=8), FGF15$^{INT\text{-}KO}$ Sham (n=8), FGF15$^{INT\text{-}KO}$ VSG (n=5). Animal number WT Sham (n=5), WT VSG (n=8), FGF15$^{INT\text{-}KO}$ Sham (n=8), FGF15$^{INT\text{-}KO}$ VSG (n=5). Data are shown as means ± S.E.M.

### Supplemental Figure 2

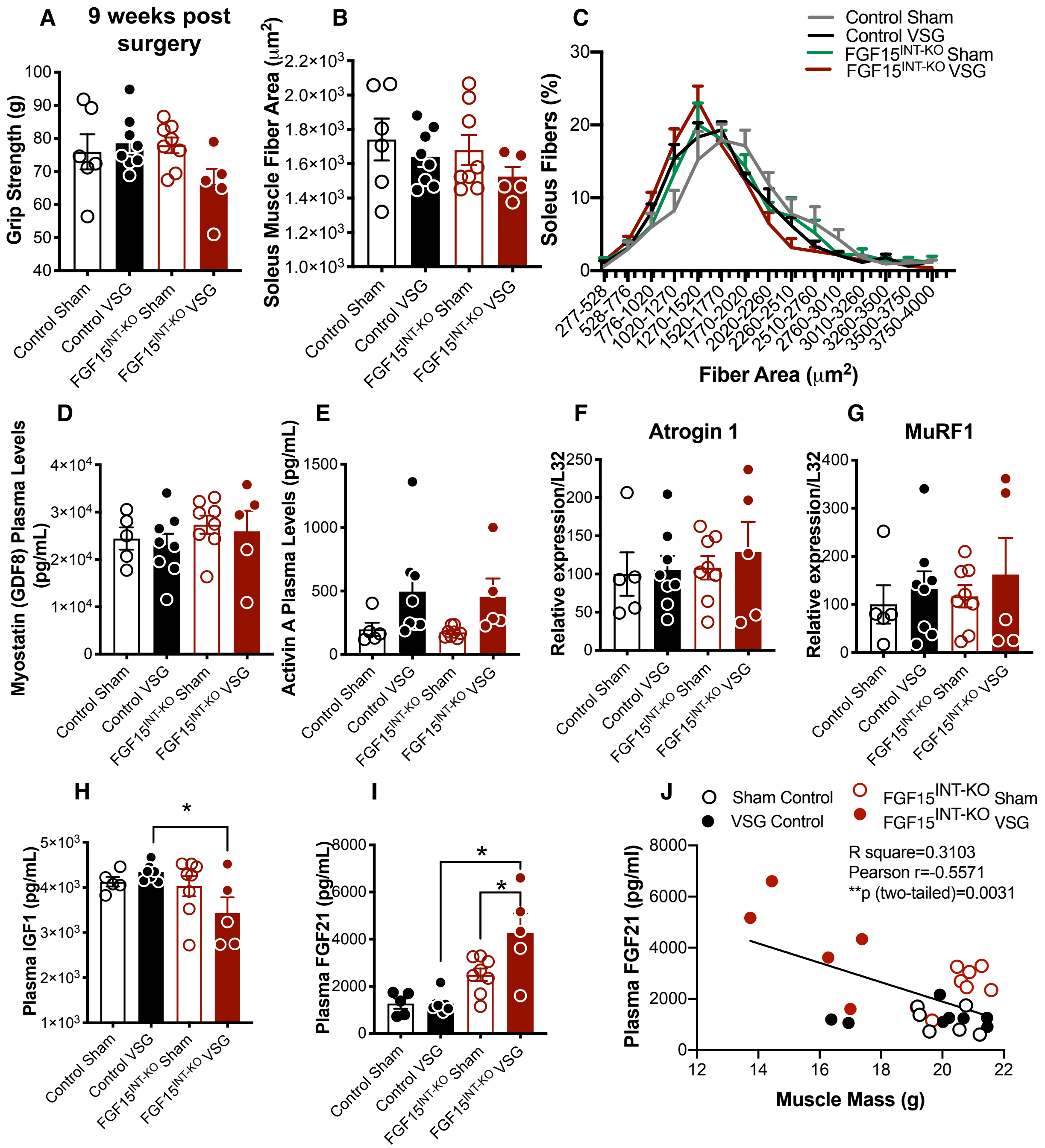

**Supplemental Figure 2. Loss in muscle mass is accompanied by decreased strength and skeletal muscle fiber size in FGF15$^{INT\text{-}KO}$ VSG mice. A.** Grip strength measured 9 weeks post-surgery. **B.** Soleus muscle average fiber area. **C.** Soleus fiber size distribution. **D.** Myostatin (GDF8) and **E.** Activin A plasma levels (postprandial, 12 weeks after surgery). RNA expression of **F.** Atrogin-1 and **G.** MuRF1 in soleus muscle. **H.** IGF-1 and **I.** FGF21 plasma levels (postprandial, 12 weeks after surgery). **J.** Correlation analysis of plasma FGF21 levels and muscle mass (12 weeks after surgery). Animal number Control Sham (n=5-6), Control VSG (n=8), FGF15$^{INT\text{-}KO}$ Sham (n=8), FGF15$^{INT\text{-}KO}$ VSG (n=5). Data are shown as means ± S.E.M. *$p<0.05$ (2-Way ANOVA with Tukey's post-test).

**Supplemental Figure 3**

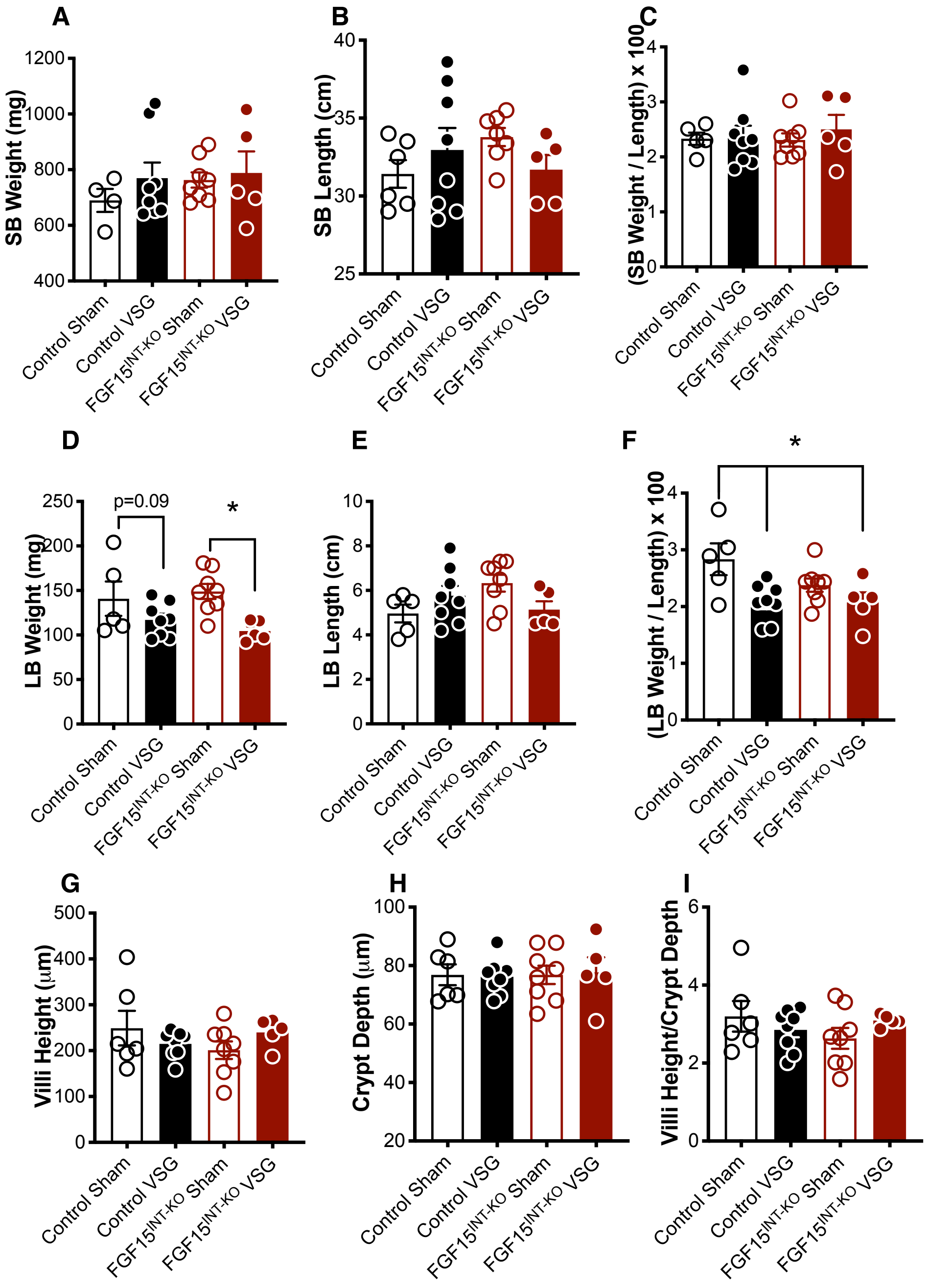


**Supplemental Figure 3. Intestinal biometry in mice lacking intestinal-derived FGF15.** SB: small bowel, LB: large bowel. **A.** SB weight. **B.** SB length. **C.** SB Weght/Length. **D.** SB weight. **E.** SB length. **F.** SB Weight/Length. **G.** Ileum villi height. **H.** Ileum crypt depth. **I.** Ileum villi height/crypt depth. Animal number Control Sham (n=5-6), Control VSG (n=8), FGF15$^{INT-KO}$ Sham (n=8), FGF15$^{INT-KO}$ VSG (n=5). Data are shown as means ± S.E.M. *p<0.05 (2-Way ANOVA with Tukey's post-test).

### Supplemental Figure 4

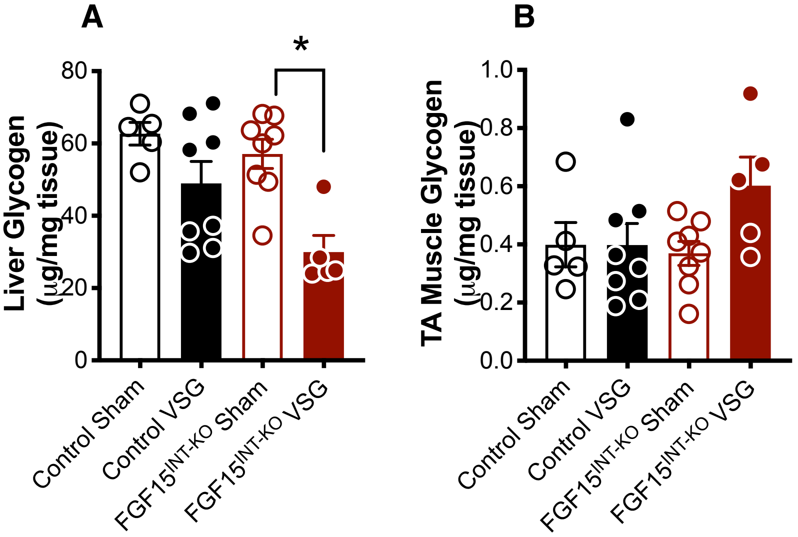


**Supplemental Figure 4. Loss of intestinal FGF15 results in aberrant glycogen metabolism following VSG. A.** Liver glycogen content and **B.** Muscle glycogen content (TA; tibialis anterior). Animal number Control Sham (n=5), Control VSG (n=8), $FGF15^{INT-KO}$ Sham (n=8), $FGF15^{INT-KO}$ VSG (n=5). Data are shown as means ± S.E.M. *$p<0.05$ (2-Way ANOVA with Tukey's post-test).

### Supplemental Figure 5

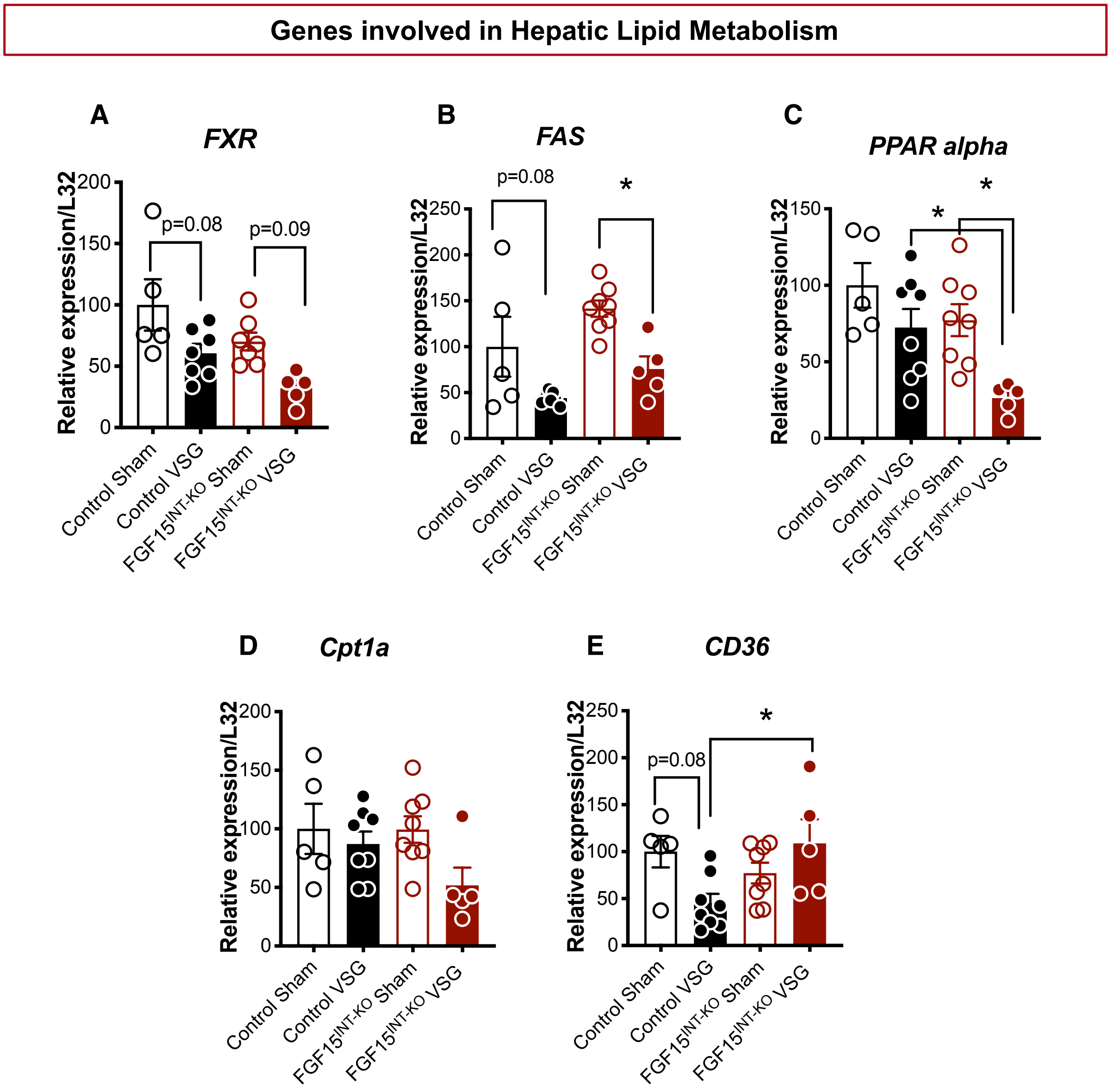


**Supplemental Figure 5. Hepatic fatty acid and lipid metabolism.** Hepatic RNA expression of **A.** Farnesoid X receptor (*FXR*) **B.** Fatty acid synthase (*FAS*) **C.** Peroxisome proliferator-activated receptor alpha (*PPAR-alpha*) **D.** Carnitine palmitoyltransferase 1A (*Cpt1a*) **E.** Cluster of differentiation 36 (*CD36*). Animal number Control Sham (n=5), Control VSG (n=8), FGF15$^{INT-KO}$ Sham (n=8), FGF15$^{INT-KO}$ VSG (n=5). Data are shown as means ± S.E.M. *p<0.05 (2-Way ANOVA with Tukey's post-test).

**Supplemental Figure 6**

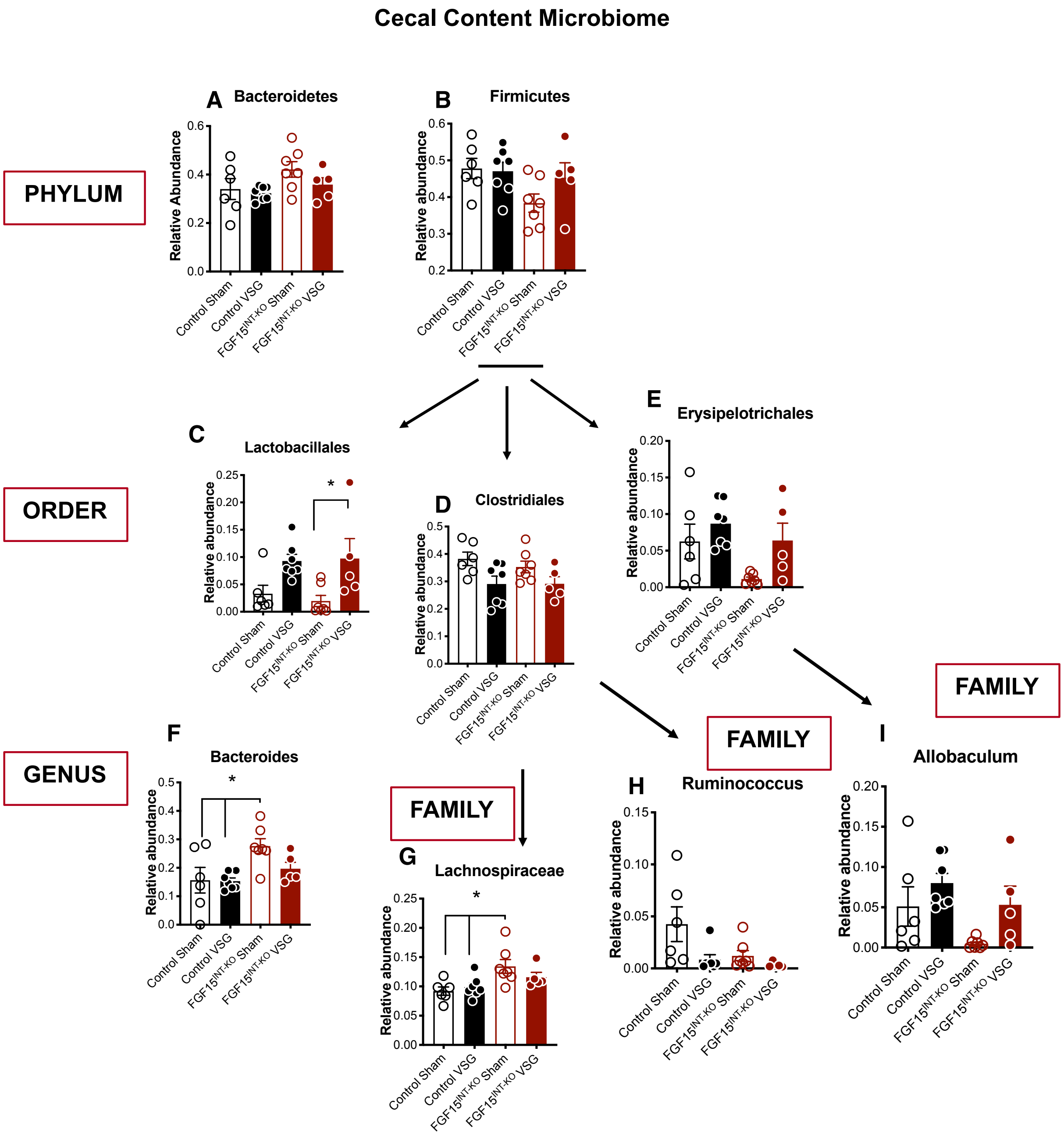


**Supplemental Figure 6. Intestinal FGF15 modulates microbiota in cecal content.** Relative abundance of **A.** *Bacteroidetes*, **B.** *Firmutes*, **C.** *Lactobacillales*, **D.** *Clostridales*, **E.** *Erysipelotrichales,* **F.** *Bacteroides*, **G.** *Lachnospiraceae*, **H.** *Ruminococcus*, **I.** *Allobaculum*. Data are shown as means ± S.E.M. *p<0.05 (2-Way ANOVA with Tukey's post-test).
